# MALT1 mediates IL-17 Neural Signaling to regulat*e C. elegans* behavior, immunity and longevity

**DOI:** 10.1101/658617

**Authors:** Sean M. Flynn, Changchun Chen, Murat Artan, Stephen Barratt, Alastair Crisp, Geoffrey M. Nelson, Sew-Yeu Peak-Chew, Farida Begum, Mark Skehel, Mario de Bono

## Abstract

Besides well-known immune roles, the evolutionarily ancient cytokine interleukin-17 (IL-17) modulates neural circuit function. We investigate how IL-17 signals in neurons, and the extent to which this signaling can alter organismal phenotypes. We combine immunoprecipitation and mass spectrometry to biochemically characterize endogenous signaling complexes that function downstream of IL-17 receptors in *C. elegans* (*Ce*) neurons. We identify the *Ce* ortholog of MALT1 as a critical output of the pathway. MALT1 was not previously implicated in IL-17 signaling or in nervous system function. MALT1 forms a complex with homologs of Act1 and IRAK and functions both as a scaffold for IκB recruitment, and as a protease. MALT1 is expressed broadly in the *Ce* nervous system, and neuronal IL-17–MALT1 signaling regulates many phenotypes, including escape behavior, associative learning, immunity and longevity. Our data suggest MALT1 has an ancient role modulating neural function downstream of IL-17 to remodel physiological and behavioral state.

---

Immune signaling pathways regulate the development and function of the nervous system in both health and disease^1–3^. Many of these effects are mediated by cytokines, small, secreted proteins that can participate in neuroimmune and inter-neuronal communication. For example, low levels of IL-1β and TNFα regulate synaptic and homeostatic plasticity in healthy animals^4, 5^; pathological levels of proinflammatory cytokines during inflammation can disrupt foetal brain development, alter adult behavior^6–9^, and drive hyperalgesia and neuroinflammatory diseases^10^. Progression of neurodegenerative diseases, including Alzheimer’s, Parkinson’s and Amyotrophic lateral sclerosis (ALS), has also been associated with chronic inflammation^11, 12^.

Recent work shows that the interleukin 17 (IL-17) pro-inflammatory cytokine can modify neural circuit activity. In a rodent model of infection during pregnancy, IL-17 secretion during maternal immune activation drives autism-related behaviors in the pups^13^. This phenotype is associated with hyperactivity of a specific cortical sub-region that expresses IL-17 receptors (IL-17R)^14^. In mice, IL-17 has also been shown to lower the activation threshold of nociceptive neurons, and contributes to mechanical hyperalgesia^15, 16^. In *C. elegans* (*Ce*) IL-17Rs are expressed throughout the nervous system, and IL-17 has been shown to act on the RMG hub interneurons, increasing their response to presynaptic input from O_2_ sensors. The increased circuit gain conferred by IL-17 ensures this animal persistently escapes 21% O_2,_ an aversive cue associated with surface exposure^17^. Specific sensory responses and behaviors are thus modulated by IL-17 across species, suggesting IL-17 has broad and conserved roles in regulating neuronal properties.

While IL-17 action on the nervous system is well established, its molecular effectors there are poorly understood. The extent to which IL-17 signaling contributes to brain function and physiology is also unclear, even in the well-defined *Ce* nervous system.

Here, we report that IL-17 signaling in the *Ce* nervous system is mediated by the paracaspase MALT-1. MALT1 has been studied extensively but almost exclusively in the mammalian immune system. It is a key signaling molecule in innate and adaptive immunity, acting downstream of ITAM- (immunoreceptor tyrosine-based activation domain) containing receptors, including the B-cell and T-cell receptors^18–20^. It has not been shown to mediate IL-17 signaling, although there has been speculation of such involvement. *In situ* hybridization suggests widespread expression of MALT1 in mouse brain, (Allen Mouse Brain Atlas, 2004), but no physiological role in neurons has been reported. We show that *Ce* MALT-1 is expressed throughout the nervous system and forms an *in vivo* complex with IL-17 signaling components, the *Ce* homologs of Act1, IRAK and IκBζ/IκBNS. We show that MALT-1 acts both as a protease and a scaffold to regulate neural function. These results represent the first described role for MALT1 signaling in the nervous system in any animal. Defects in IL-17/MALT-1 signaling lead to reconfigured gene expression, and altered behavior and physiology, including enhanced immunity and extended lifespan.

## Results

### Proteomic analysis of IL-17 signaling identifies an ACTL-1–IRAK–MALT-1– NFKI-1 complex in *C. elegans*

IL-17 appears to act mostly on the nervous system in *Ce*^17^. We studied this signaling using proteomics. We epitope tagged all soluble IL-17 pathway components highlighted by genetics^17^, immunoprecipitated (IP’d) them from *Ce* extracts, and identified interacting proteins using mass spectrometry (MS, Fig. 1a).

**Figure 1.**
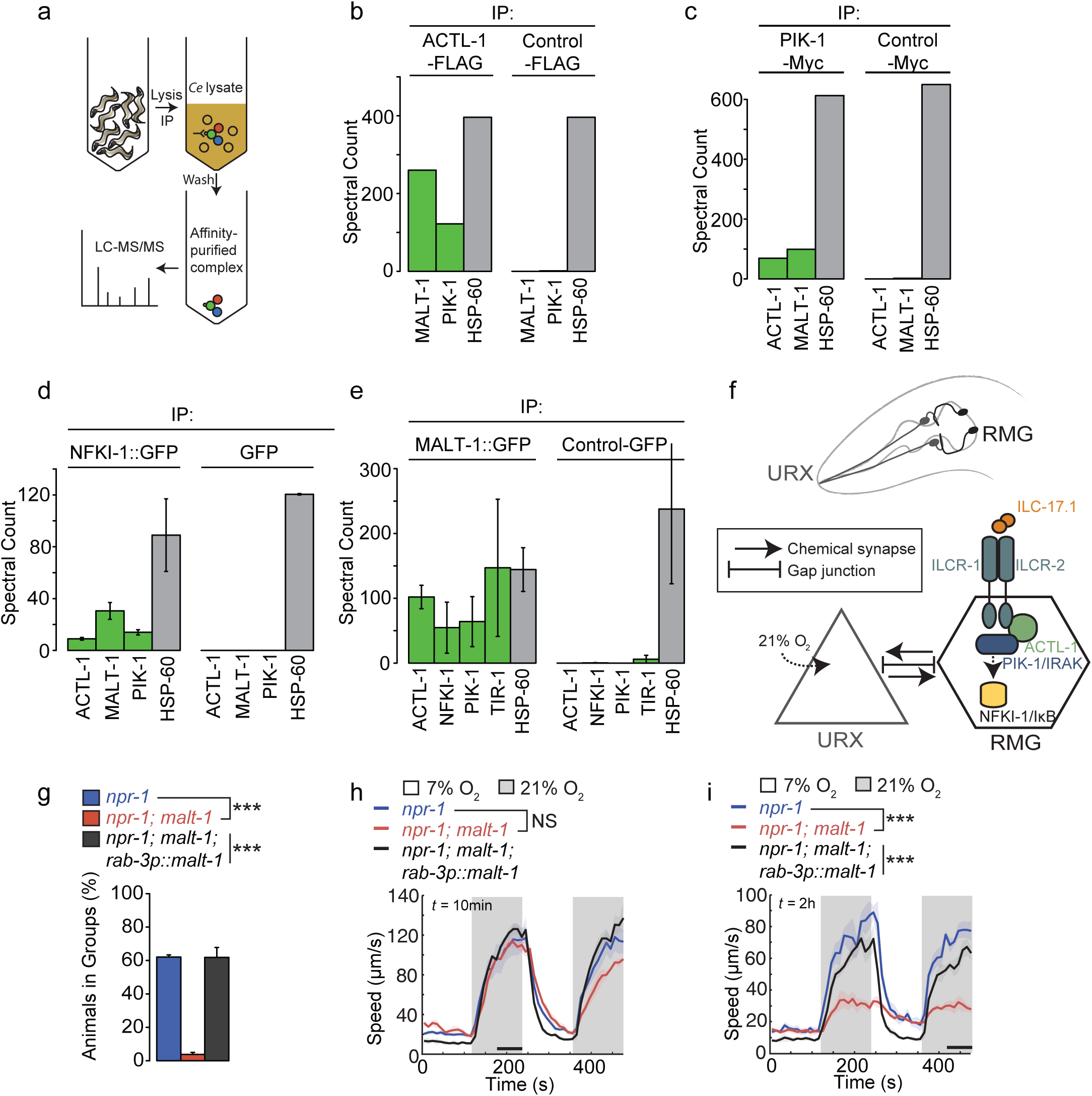
MALT-1 forms a complex with ACTL-1, PIK-1/IRAK and NFKI-1. **a** Schematic for affinity-purification and LC-MS/MS analysis of epitope-tagged IL-17 signaling components from *C. elegans* extracts. **b**-**e** Pull-down of ACTL-1-FLAG, PIK-1-Myc or NFKI-1::GFP specifically co-IPs MALT-1 (**b**-**d**). Conversely, pull-down of MALT-1::GFP specifically co-IPs ACTL-1, PIK-1 and NFKI-1 (E). Bars show total spectral counts, a semiquantitative readout of abundance^64^, and error bars represent SEM, N=2 (**e**), N=3 (**f**). Grey bars show levels of HSP-60, a common contaminant. **f** Schematic of IL-17 signaling in O_2_-escape circuit. Upshifts in [O_2_] are sensed by URX neurons, which tonically signal to RMG hub interneurons. IL-17 signaling increases the responsiveness of RMG neurons to promote escape from 21% O_2_. **g** *malt-1* promotes *Ce* aggregation (N ≥ 4 assays). ***, P < 0.001, ANOVA with Tukey’s post hoc HSD. **h** and **i** *malt-1* mutants are strongly aroused by 21% O_2_ if stimulated immediately after transfer to the assay plate (**h**), but respond weakly to 21% O_2_ if allowed to settle over a 2h period (**i**), n ≥ 46 animals. Plots show average speed (line) and SEM (shaded regions). Time of assay after transfer is shown at top left. ***, P < 0.001, Mann-Whitney *U* test (speed). Here and in subsequent figures, black bars indicate time intervals used for statistical comparisons. See also Supplementary Figures 1-4 and Supplementary Table 1.

ACTL-1 and PIK-1 are *Ce* orthologs of mammalian Act1 and IRAKs respectively, and signal downstream of the *Ce* IL-17 co-receptors ILCR-1 and ILCR-2^17^. Genetic analysis suggests NFKI-1, a homolog of mammalian I*κ*B*ζ* and I*κ*BNS, acts downstream of ACTL-1, PIK-1 and the IL-17Rs^17^.

We tagged endogenous ACTL-1 with a FLAG epitope, endogenous PIK-1 with a Myc epitope, and integrated an *nfki-1::gfp* transgene. We showed the tagged proteins were functional (Supplementary Fig. 1), and then IP’d them from *Ce* extracts. As controls, we also IP’d proteins unrelated to IL-17 signaling tagged with the same epitopes. Using mass spectrometry (LC-MS/MS) we then identified specific interactors for each signaling component (Fig. 1b-d and Supplementary Table 1a-c).

As expected from non-physiological co-IP experiments using mammalian tissue culture cells^17^, PIK-1 co-precipitated specifically with ACTL-1 (Fig. 1b), and reciprocally, ACTL-1 co-precipitated specifically with PIK-1 (Fig. 1c). IP of NFKI-1 also identified ACTL-1 and PIK-1/IRAK as specific interactors, suggesting these proteins form a complex *in vivo* (Fig. 1d). We identified many other apparently specific interactors for each component. These are listed in Supplementary Table 1 as a resource.

One interactor, the *Ce* ortholog of the paracaspase MALT1, consistently co-IP’d with each of ACTL-1, PIK-1 and NFKI-1 (Fig. 1b-d). MALT1 paracaspases are cysteine proteases with specificity for arginine residues^21, 22^. Their caspase-like protease domain is highly conserved, as is their domain organization, consisting of an N-terminal death domain (DD) followed by 2-3 Ig (immunoglobulin)-like motifs that flank the paracaspase domain (Supplementary Fig. 2a)^23^. Mammalian MALT1 signals downstream of B cell, T cell, and other cell surface receptors containing an ITAM motif, forming a filamentous complex called the CBM signalosome that contains a CARD domain protein, BCL10, and MALT1^18–20^ (Supplementary Fig. 2b). MALT1 functions in the immune system are under intense scrutiny, but its roles elsewhere, and in invertebrates, have not been established.

To confirm these biochemical interactions we expressed functional, GFP-tagged MALT1 pan-neuronally, and identified interacting partners using IP/MS of extracts from the transgenic *Ce* strain. As a control, we performed IP/MS on extracts from strains expressing GFP-tagged neuronal proteins unrelated to IL-17 signaling. ACTL-1, PIK-1 and NFKI-1 each interacted specifically with MALT-1-GFP (Fig. 1e). We also identified other specific interactors (Supplementary Table 1d) including the *Ce* ortholog of mammalian SARM1, called TIR-1, which is implicated in the immune response^24, 25^, left/right asymmetry of an olfactory neuron^26^, and experience-dependent plasticity^27^. MALT-1 also interacted specifically with a large group of proteins implicated in RNA metabolism, including splicing factors and polyA binding proteins, suggesting it may localize to the nucleus or ribonucleoprotein particles (RNPs) (Supplementary Fig. 3).

### MALT-1 promotes *C. elegans* aggregation and escape from 21% O_2_

Previous work has not implicated MALT1 in IL-17 signaling or neural function. In *Ce*, the IL-17 ILC-17.1 signals through the ILCR-1/ILCR-2 receptors on the RMG interneurons to increase RMG responsiveness to input from their pre-synaptic partner, the URX O_2_-sensing neurons (Fig. 1f). Increased RMG signaling enables *Ce* to strongly and persistently escape 21% O_2_ and to aggregate^17, 28, 29^. To probe the functional relevance of our proteomics data we asked if the large collection of mutants that yielded IL-17 pathway mutants^17^ included *malt-1* loss-of-function alleles. This collection was isolated in a forward genetic screen for mutants defective in escape from 21% O_2_ and aggregation behavior. Four strains in the collection harbored *malt-1* alleles; one introduced a premature stop codon; another mutated the highly conserved E464 residue (Supplementary Fig. 4a and b), known to be essential for catalytic activity in mammalian MALT1^30^. We mapped the aggregation defect of this strain to an interval containing *malt-1* (Supplementary Fig. 4c). Targeted disruption of *malt-1* using CRISPR/Cas9 resulted in an aggregation-defective strain whose phenotype could be rescued using a wild-type *malt-1* transgene (Fig. 1g; and Supplementary Fig. 4e). These data confirm that MALT-1, like IL-17 signaling, promotes aggregation.

*C. elegans* aggregate to escape 21% O_2_, which signals surface exposure^31–33^. 21% O_2_ evokes sustained arousal in wild *Ce* isolates^34^, and in *npr-1 (n*euro*p*eptide *r*eceptor *1)* mutants of the domesticated N2 lab strain^35^. *npr-1* mutants defective in IL-17 signaling are not aroused by 21% O_2_ in the absence of additional recent mechanosensory stimulation. Moreover, the arousal evoked when such animals are stimulated by 21% O_2_ shortly after being mechanically disturbed (e.g by being picked) is not sustained^17^. *malt-1* mutants showed this hallmark phenotype (Fig. 1h, i), consistent with MALT-1 playing a role in *Ce* IL-17 signaling.

### MALT-1 modulates responsiveness of RMG interneurons

*malt-1::GFP* and *malt-1::RFP* transgenes were expressed broadly in the nervous system (Fig. 2), including in the O_2_-sensing neurons AQR, PQR and URX (Supplementary Fig. 5a and b) and their post-synaptic partner the RMG interneurons (Fig. 3a). *malt-1* phenotypes were rescued by expressing *malt-1* cDNA pan-neuronally, confirming that MALT-1 has neuronal functions (Fig. 1g, i). Selectively expressing *malt-1* cDNA in the RMG interneurons, or the O_2_-sensing neurons restored aggregation to *malt-1* mutants (Fig. 3b), but only partially rescued the O_2_-response defects (Fig. 3c). By contrast, we observed almost complete rescue when we expressed MALT-1 simultaneously in both sets of neurons (Fig. 3c; Supplementary Fig. 4e and f). Thus, like IL-17Rs^17^, MALT-1 functions in RMG and AQR, PQR and URX to promote escape from 21% O_2_.

**Figure 2.**
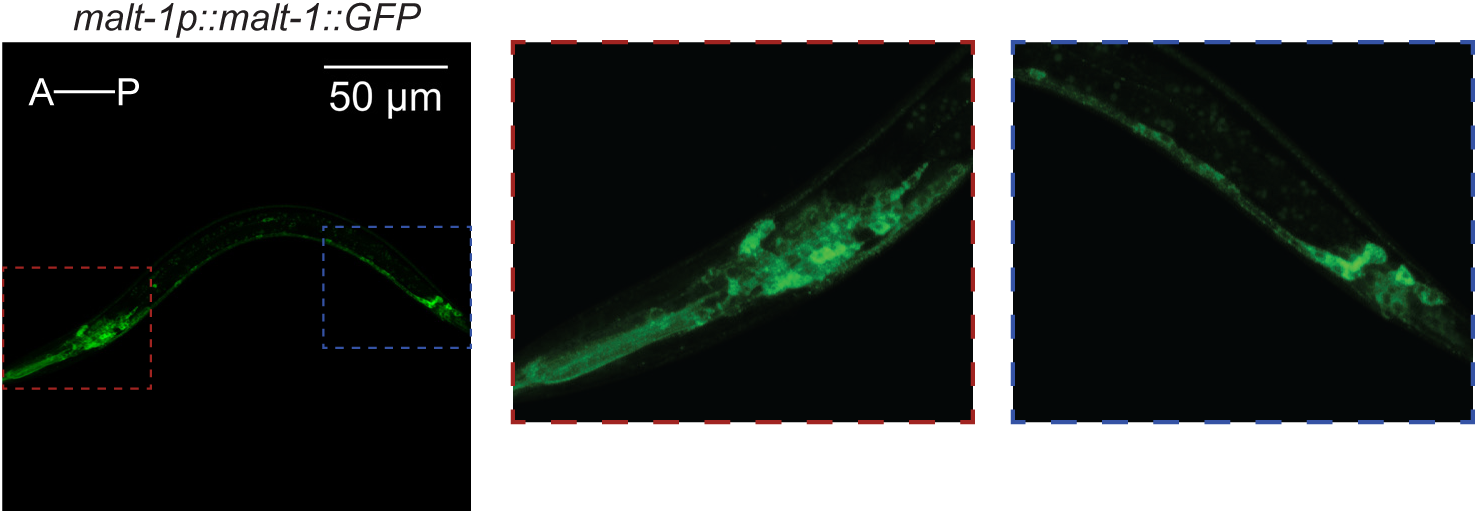
MALT-1 is expressed widely in the nervous system. A transgene expressing C-terminally GFP-tagged MALT-1 from its endogenous promoter (4kb of upstream DNA) is expressed broadly in the nervous system, including many neurons in the head (red box) and tail (blue box). See also Supplementary Figure 5.

**Figure 3.**
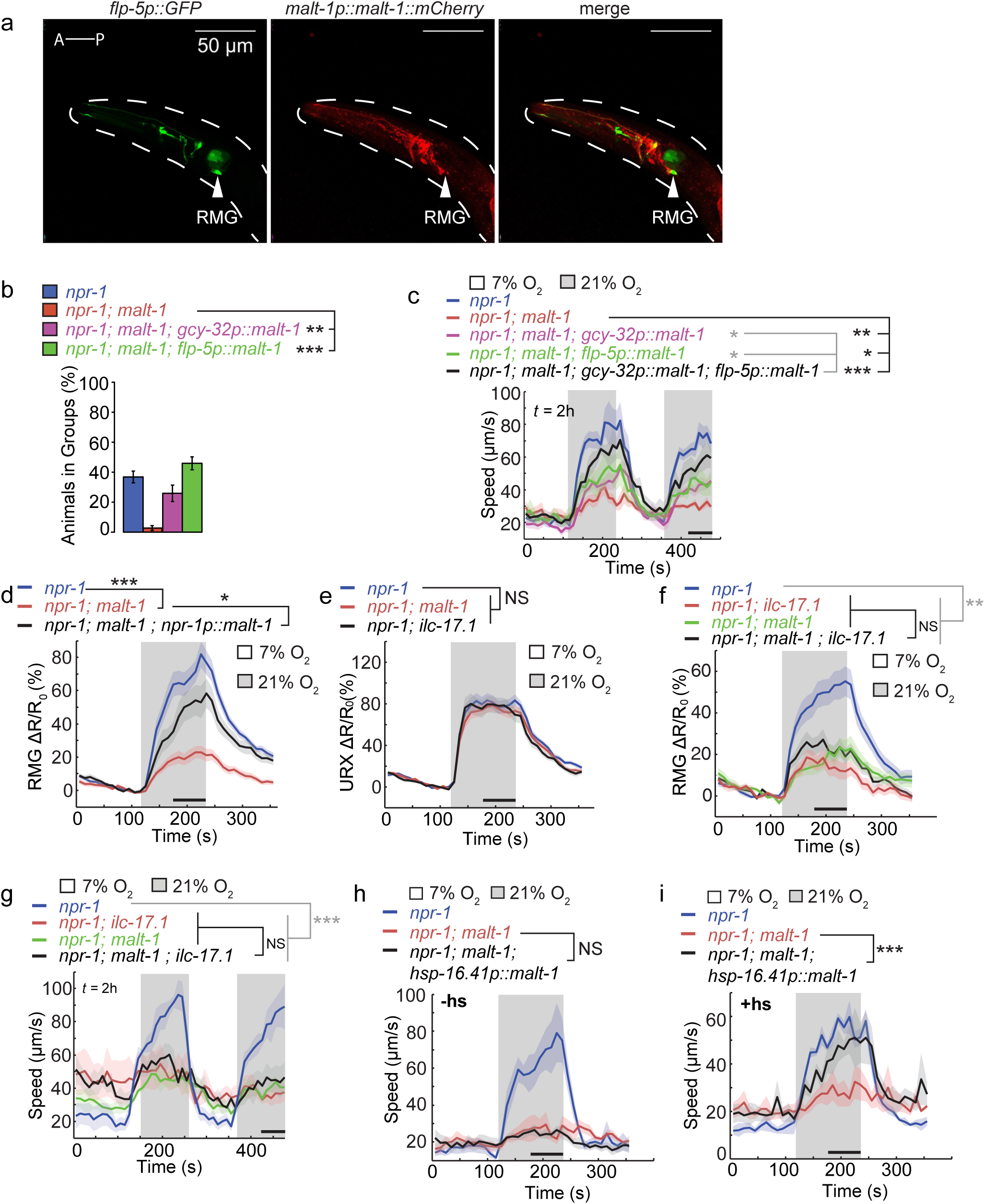
MALT-1 mediates IL-17 signaling in RMG interneurons. **a** A MALT-1::mCherry translational fusion, expressed from its endogenous promoter (4kb), is expressed in RMG interneurons. RMG is recognized by its characteristic shape, location, and using a *flp-5p::gfp* reporter (**c**, Kim and Li, 2004). **b** Expressing *malt-1* cDNA from either the *flp-5* promoter (RMG, ASG, PVT, I4, M4 and pharyngeal muscle), or the *gcy-32* promoter (URX, AQR and PQR) rescues the aggregation defect of *malt-1* mutants. N = 4 assays. **, P < 0.01, ***, P < 0.001, ANOVA with Tukey’s post hoc HSD. **c** The O_2_-response defect of *malt-1* mutants is partially rescued by expressing *malt-1* cDNA from the *flp-5* promoter (RMG, ASG, PVT, I4, M4 and pharyngeal muscle), or the *gcy-32* promoter (URX, AQR and PQR), and almost completely rescued when *malt-1* is expressed from both promoters simultaneously. Lines indicate average speed and shaded regions indicate SEM. n ≥ 46 animals. *, P < 0.05, **, P < 0.01, ***, P < 0.001, Mann-Whitney *U* test. **d** and **e** Disrupting *malt-1* attenuates Ca^2+^ responses evoked by 21% O_2_ in RMG (D, n ≥ 25 animals) but not URX (**e**, n ≥ 13 animals) neurons. The RMG defect can be rescued by expressing *malt-1* cDNA in both RMG and URX, using the *npr-1* promoter (**d**). Ca^2+^ responses are reported by YC2.60 cameleon. *, P < 0.05, ***, P < 0.001, Mann-Whitney *U* test. **f** and **g** Null mutations in *malt-1* and *ilc-17.1* do not show additive phenotypes when either RMG Ca^2+^ transients (**f,** n ≥ 24 animals) or speed responses evoked by 21% O_2_ are measured (**g**, n ≥ 50 animals). *, P < 0.05, **, P < 0.01, Mann-Whitney *U* test. **h** A transgene expressing *malt-1* cDNA from the *hsp-16.41* promoter does not rescue *malt-1* phenotypes in the absence of heat-shock (n ≥ 33 animals). **i** Heat-shock-induced cDNA expression in adults restores O_2_–evoked responses to *malt-1* mutants. Plots show average speed (line) and SEM (shaded regions); n ≥ 35 animals. ***, P < 0.001, Mann-Whitney *U* test.

Ca^2+^ imaging revealed that O_2_-evoked Ca^2+^ responses in RMG were significantly reduced in *malt-1* mutants, both in freely moving (Supplementary Fig. 4g) and immobilized animals (Fig. 3d). By contrast, O_2_-evoked Ca^2+^ responses in the URX sensory neurons appeared normal in *malt-1* mutants (Fig. 3e). These phenotypes recapitulate those observed in IL-17 signaling mutants^17^. The RMG Ca^2+^ response defect was rescued by expressing *malt-1* cDNA from the *npr-1* promoter, which drives expression in RMG and the AQR, PQR and URX neurons (Fig. 3d).

The *malt-1* and *ilc-17.1* mutant phenotypes were not additive. Both the Ca^2+^ signaling (Fig. 3f) and behavioral response (Fig. 3g) defects of *malt-1; ilc-17.1* double mutants resembled those of single mutants, suggesting MALT-1 and ILC-17.1 function in the same pathway. Similarly, the RMG response defects of *malt-1* mutants were not enhanced by defects in PIK-1/IRAK (Supplementary Fig. 4g).

Together, our biochemical, genetic, behavioral and physiological data suggest that the paracaspase MALT-1 mediates IL-17 signaling in neurons, most likely via a signaling complex made up of ACTL-1–IRAK/PIK-1–MALT-1–NFKI-1. To examine if *malt-1* is required developmentally, we expressed it selectively in adults using a heat-shock-inducible promoter. Without heat-shock, the *phsp-16::malt-1* cDNA transgene did not rescue the O_2_-response phenotype of *malt-1* mutants (Fig. 3h). Heat-shock-induced expression during the 4^th^ larval stage was sufficient to restore behavioral responses (Fig. 3i), suggesting that MALT-1, like other IL-17 signaling components^17^, can alter circuit properties after the circuits have developed.

### MALT-1 functions as a protease in the nervous system

In the mammalian immune system MALT1 functions both as a scaffold and as a protease. To examine if MALT-1 acts as a protease in neurons we mutated the active site cysteine of the endogeneous *malt-1* gene to alanine. The equivalent mutation is used in a paracaspase-dead model in mice^36–38^. *malt-1 C374A* animals resembled *malt-1* null mutants, and could be rescued by pan-neuronal expression of *malt-1* cDNA (Fig. 4a; Supplementary Fig. 6a). By contrast, a *malt-1 C374A* transgene was unable to rescue the phenotype of *malt-1(db1194)* mutants (Fig. 4b). Unexpectedly, overexpressing *malt-1 C374A* in a WT background conferred a *malt-1(null)* phenotype (Fig. 4c), suggesting that catalytically dead MALT-1 can act as a dominant negative. Together these data suggest that MALT-1 signals as a protease in the *C. elegans* nervous system.

**Figure 4.**
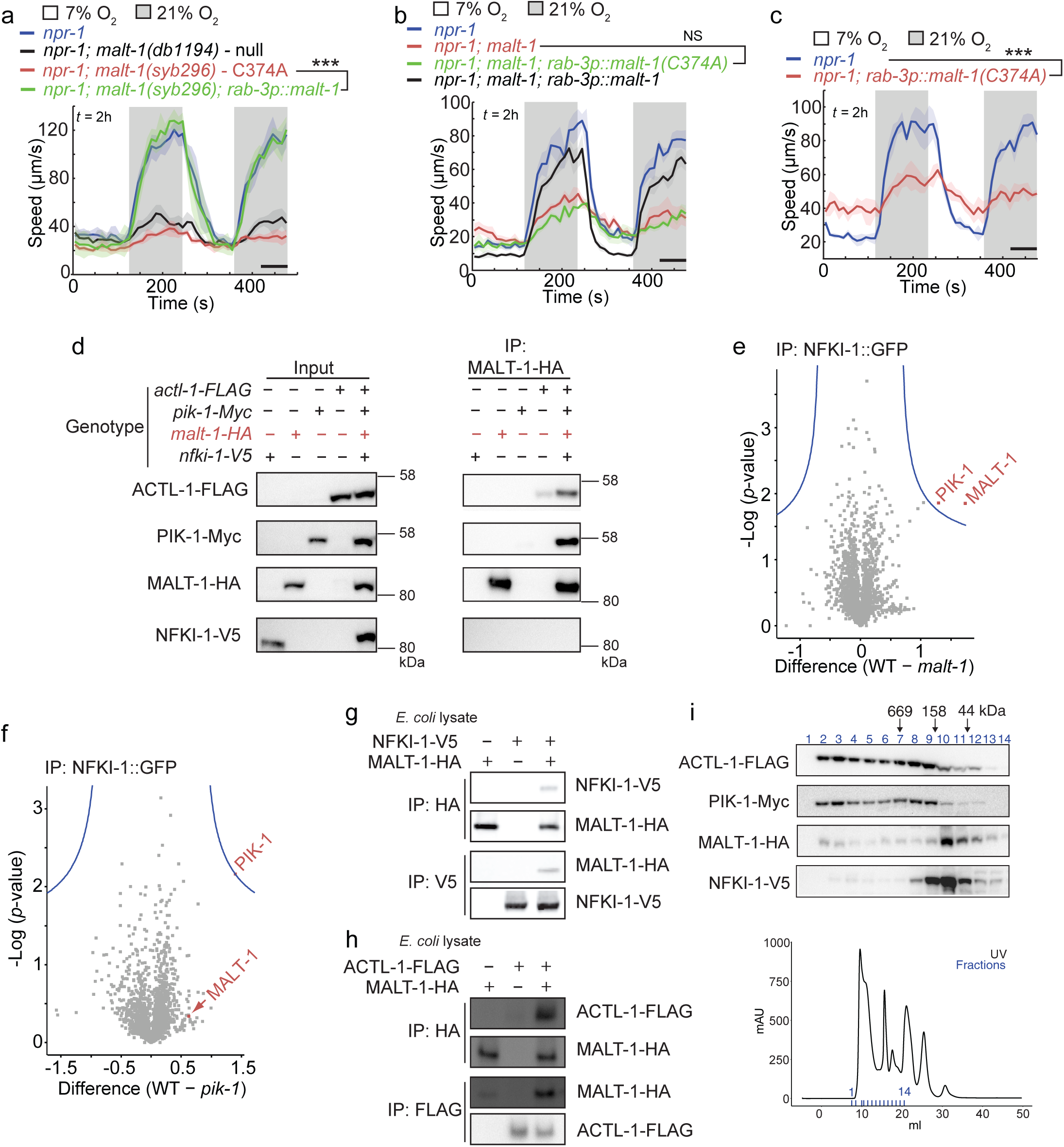
MALT-1 has scaffolding and enzymatic roles in IL-17 signaling. **a-c** MALT-1’s function in the nervous system requires its proteolytic activity. (**a**) *malt-1(syb296)* mutants that express a catalytically inactive MALT-1 (C374A) show O_2_ response defects comparable to those of *malt-1(null)* animals. Pan-neuronal expression of *malt-1* cDNA rescues this phenotype (n ≥ 29 animals). ***, P < 0.001, Mann-Whitney *U* test. **b** cDNA encoding a MALT-1 C374A catalytically inactive protein, expressed from the *rab-3* promoter, does not rescue the O_2_ response defects of *malt-1* mutants (n ≥ 46 animals). Data corresponding to *npr-1* and *npr-1; malt-1; rab-3p::malt-1* in **b** are the same as those shown in Fig. 1i, and were obtained in parallel to the genotypes shown. ***, P < 0.001, Mann-Whitney *U* test. **c** Overexpressing MALT-1 C374A cDNA in *npr-1* animals inhibits the arousal response to 21% O_2_ (n ≥ 53 animals). ***, P < 0.001, Mann-Whitney *U* test. **d** Endogenous ACTL-1 and PIK-1 co-IP with endogenous MALT-1 in *npr-1* and *npr-1; ilc-17.1* animals. Endogenous NFKI-1 was not detected. Tags were knocked in by CRISPR. **e** and **f** Volcano plot showing quantitative LC-MS/MS of proteins that interact with NFKI-1::GFP in *malt-1* and *pik-1* mutants compared to wild type. NFKI-1::GFP was purified using GFP-Trap beads, and immunoprecipitated proteins labeled using tandem mass tags (TMT-labelling). The average relative abundance in two biological replicates is shown. Blue lines indicate a significance threshold of P= 0.05 (two sample t-test). The amount of PIK-1 that co-IPs with overexpressed NFKI-1::GFP is significantly reduced in *malt-1* mutants (**e**). The relative amount of MALT-1 that co-IPs with NFKI-1 is not significantly decreased in *pik-1* mutants (**f**). **g** and **h** IPs of His10-tagged *Ce* ACTL-1-FLAG, MALT-1-HA, and NFKI-1-V5 recombinantly expressed in *E. coli* show that MALT-1 can directly bind NFKI-1 (**g**) and ACTL-1 (**h**). **i** Elution profiles of ACTL-1, PIK-1, MALT-1 and NFKI-1 proteins in a *C. elegans* extract run on a Superose 6 Gel Filtration column and visualized by immunoblot. All four proteins can be found in very high molecular weight complexes. See also Supplementary Figure 6.

We also asked if IL-17 signaling requires PIK-1/IRAK kinase activity. We created a single copy transgene in which the ATP-binding pocket lysine residue (K217) was mutated to alanine. The *K217A* transgene rescued *pik-1(null)* phenotypes (Supplementary Fig. 6b), suggesting that kinase activity is not essential for PIK-1 to regulate behavior.

### MALT-1 promotes assembly of IL-17 signaling complexes

To extend our *in vivo* proteomic analyses we made a strain in which endogenous ACTL-1, PIK-1, MALT-1 and NFKI-1 were each tagged with different epitopes. To corroborate our LC-MS/MS data we first showed that ACTL-1 and PIK-1 specifically co-IP’d with MALT-1 (Fig. 4d). When we IP’d ACTL-1-FLAG and PIK-1-Myc from our knock-in strains we did not detect NFKI-1 by mass spectrometry (Supplementary Table 1a, b). We also failed to detect NFKI-1-V5 by Western blot when we IP’d MALT-1-HA (Fig. 4d) from a multiple knock-in strain. These results suggest the association of NFKI-1 with the complex may be transient or labile under basal conditions, and only detectable when NFKI-1 or MALT-1 is overexpressed (Fig. 1d, e).

To analyse the signaling complex further we carried out IPs from strains overexpressing NFKI-1-GFP. When we quantitatively compared NFKI-1 complexes from WT, *malt-1* and *pik-1* mutants, using IP/MS, we found that the amount of PIK-1/IRAK co-precipitating with NFKI-1 was reduced when MALT-1 was absent (Fig. 4e). By contrast, in *pik-1* mutants the interaction between MALT-1 and NFKI-1 was not significantly reduced (Fig. 4f). These data suggest that NFKI-1 is recruited to signaling complexes by MALT-1.

To ask if MALT-1 and NFKI-1 interact directly, we expressed epitope-tagged versions of the proteins in *E. coli*, and performed pairwise tests for co-immunoprecipitation. MALT-1-HA IP’d NFKI-1-V5, and conversely NFKI-1-V5 IP’d MALT-1, supporting a direct physical interaction (Fig. 4g). MALT-1 also interacted directly with ACTL-1 (Fig. 4h). These data suggest that MALT-1 provides a bridge between IL-17R associated complexes and NFKI-1.

### Sub-cellular localization of IL-17 signaling components

In the mammalian immune system IRAKs and MALT1 are core components of the Myddosome and CBM signalosome, respectively. These complexes are structurally related filamentous oligomers that assemble in the cytosol^39, 40^. IκB family proteins perform both cytoplasmic and nuclear functions downstream of signalosome assembly^41^. Fractionation of a *Ce* lysate by gel filtration revealed that ACTL-1 and PIK-1 exist mostly as high-molecular weight species; they eluted in the heaviest fractions, including the void, of a gel filtration column (Fig. 4i). MALT-1 and NFKI-1 ran mostly as smaller species (∼50-200kDa), but they were also detectable in the heavier ACTL-1- and PIK-1-containing fractions. These observations suggest that a ternary complex, potentially related to the Myddosome and the CBM signalosome but incorporating ACTL-1/PIK-1/MALT-1, may mediate IL-17 signaling in *Ce* neurons.

To determine the sub-cellular localization of IL-17 signaling components, we separated the nuclear and cytosolic fractions of our lysate. ACTL-1-FLAG, PIK-1-Myc and MALT-1-HA were detected in both cytoplasmic and nuclear fractions (Fig. 5a), while NFKI-1-V5 was predominantly in nuclear fractions (Fig. 5a). It is not clear what function IL-17 signaling proteins serve in the nucleus, however it is notable that NFKI-1 specifically co-IP’d chromatin state modifiers, including CREB binding protein (CBP), a histone acetyltransferase^42^, suggesting that NFKI-1 regulates transcription (Supplementary Table 1c).

**Figure 5.**
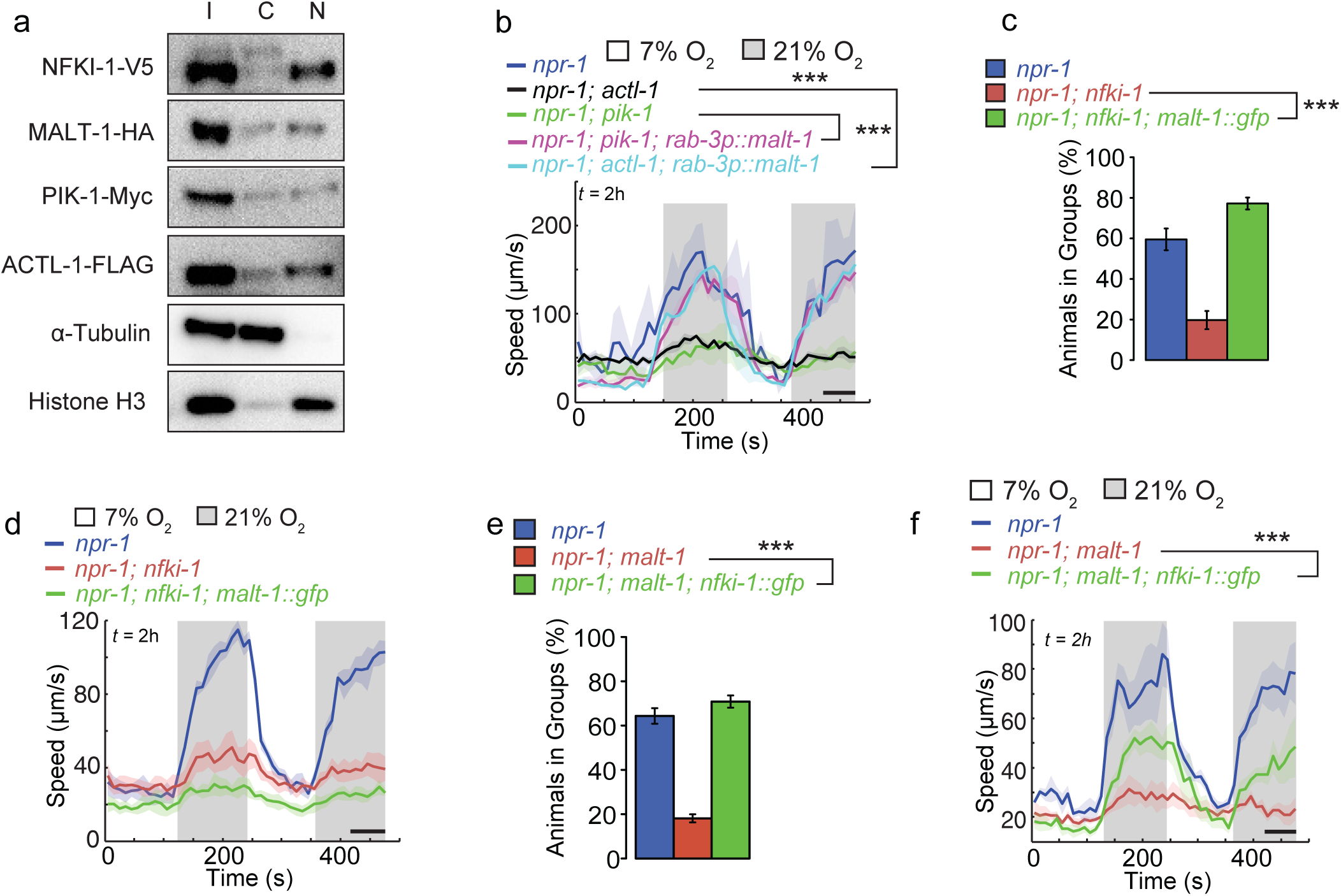
MALT-1 and NFKI-1 provide partially parallel outputs of IL-17 signaling. **a** Immunoblot analysis of IL-17 signaling components from nuclear and cytoplasmic fractions of *Ce* lysate. I,input, C, cytosolic, N, nuclear. NFKI-1 is predominately nuclear; ACTL-1, PIK-1 and MALT-1 are distributed between the nucleus and cytoplasm. **b** Overexpressing *malt-1* gDNA restores the arousal response to 21% O_2_ to *actl-1* and *pik-1* mutants (n ≥ 19 animals). **c** and **d** Overexpressing *malt-1* gDNA also rescues the aggregation phenotype (**c**, N ≥ 6 assays), but not the arousal defect of *nfki-1* mutants (**d**, n ≥ 39 animals). (**e** and **f**) The aggregation phenotype of *malt-1* is rescued by overexpressing *nfki-1* cDNA (**e**, N ≥ 4 assays), while speed defects are partially rescued (**f**, n ≥ 36 animals). ***, P < 0.001, Mann-Whitney *U* test (speed) or ANOVA with Tukey’s post hoc HSD (aggregation).

### MALT-1 and NFKI-1 provide parallel outputs for IL-17 signaling

Overexpressing NFKI-1 suppresses *actl-1* and *pik-1* null phenotypes, suggesting NFKI-1 functions downstream of those signaling components^17^. Overexpressing MALT-1 also rescued the O_2_ arousal defects of *actl-1* and *pik-1* mutants (Fig. 5b). To test whether MALT-1 functions upstream or downstream of NFKI-1, we asked whether overexpressing either component rescued a null mutant of the other. Overexpressing NFKI-1 in *malt-1(null)* mutants, or MALT-1 in *nfki-1(null)* animals, fully rescued the aggregation defect but either did not restore, or only partly restored, the arousal response to 21% O_2_ (Fig. 5c-f). These data suggest MALT-1 and NFKI-1 provide partially parallel outputs for IL-17 signaling.

### Disrupting IL-17 signaling reprograms gene expression

In mammalian tissues IL-17 acts globally to drive pro-inflammatory gene expression^43^. We defined a transcriptional fingerprint of *Ce* IL-17 signaling by comparing the whole-animal RNA-seq profiles of *ilc-17.1*, *malt-1* and *nfki-1* mutants to that of controls (Supplementary Table 2). Data analysis suggested that pathways implicated in neuropeptide signaling, metabolism, ageing, and immunity were significantly altered by IL-17 signaling (Fig. 6a and b, and Supplementary Table 2).

**Figure 6.**
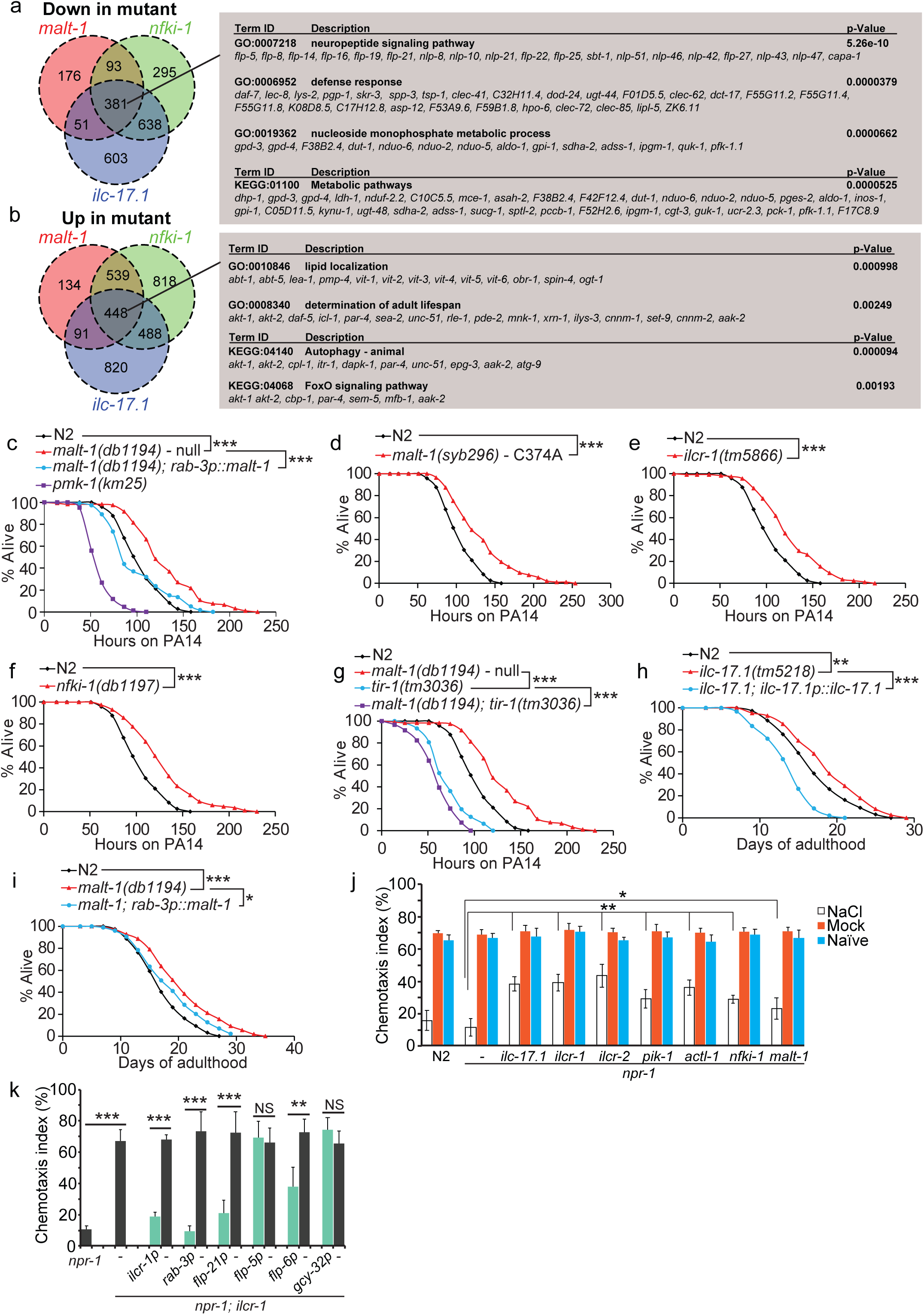
MALT-1 acts downstream of IL-17 signaling to reprogram behavior and physiology. **a** and **b** Summary of number of genes that are significantly downregulated (**a**) and upregulated (**b**) in whole animal RNA-seq profiles of *malt-1; npr-1, ilc-17.1; npr-1* and *nfki-1; npr-1* mutants compared to *npr-1*. The gene ontology (GO) terms and KEGG pathways significantly overrepresented among genes dysregulated in all three mutant conditions are shown. All changes shown are statistically significant (q-value < 0.05), with a minimum log2(fold-change) of ±0.25. **c**-**f** Mutants defective in *malt-1* and other IL-17 signaling components are resistant to *P. aeruginosa* PA14. The enhanced survival of *malt-1* mutants compared to N2 controls is rescued by pan-neuronal expression of *malt-1* gDNA. *, P, < 0.05, **, P, < 0.01, ***, P, < 0.001, logrank test (n=120 animals). **g** The enhanced resistance of *malt-1* mutants to PA14 requires TIR-1. Like *tir-1* mutants, *malt-1; tir-1* double mutants are hypersensitive to PA14 infection. ***, P < 0.001, logrank test (n=120 animals). **h** and **i** The lifespan of *ilc-17.1* and *malt-1* mutants is increased compared to N2 controls. The *ilc-17.1* phenotype can be rescued by expressing *ilc-17.1* cDNA from its endogenous promoter, and the *malt-1* phenotype is rescued by expressing *malt-1* gDNA pan-neuronally. **, P < 0.01, ***, P < 0.001, logrank test (n=180 animals). **j** and **k** Salt chemotaxis after conditioning by food-withdrawal in the absence or presence of NaCl. **j** *ilc-17.1*, *ilcr-1, ilcr-2*, *pik-1*, *actl-1*, *nfki-1* and *malt-1* mutants show a NaCl association learning defect: they retain stronger chemotaxis to NaCl after conditioning than wild-type controls. **k** The salt chemotaxis learning defect of *ilcr-1* mutants is rescued by driving *ilcr-1* expression in many neurons (*rab-3* or *flp-21* promoters), or specifically in ASE (*flp-6* promoter). For simplicity, only the chemotaxis index of animals previously conditioned with NaCl in the absence of food is shown. For each transgenic strain we compare the behavior of animals bearing the transgene to their non-transgenic siblings. *, P < 0.05, **, P < 0.01, ***, P < 0.001, ANOVA with Tukey’s post hoc HSD, N ≥ 6 assays. See also Supplementary Figure 7 and Supplementary Tables 2, 3, 4 and 5.

**Figure 7.**
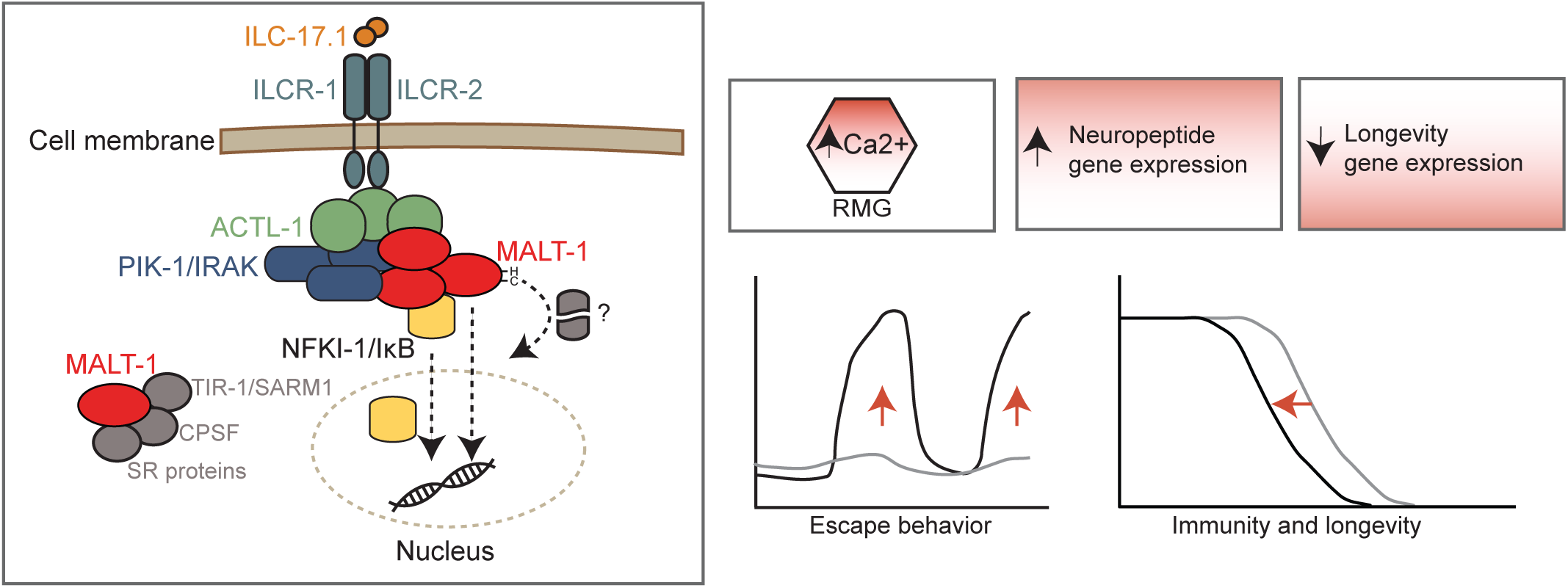
Model. Activation of nematode IL-17Rs engages ACTL-1, the *Ce* ACT1-like adaptor, probably via their SEFIR domains. ACTL-1 recruits the *Ce* IRAK and MALT1 homologs to form the ACT1-IRAK-MALT1 signalosome in the cytoplasm. This serves a scaffolding function to recruit IκBζ/NFKI-1, and modulate its actvity by an unknown mechanism. IκBζ, probably orchestrates changes in the transcriptome of RMG and other cells that result in increased neuronal output by cooperating with other transcriptional regulators in the nucleus,. MALT1-mediated cleavage of unknown substrate(s) positively regulates IκBζ signaling. In parallel to this pathway, MALT-1 forms a complex with TIR-1/SARM1 to promote PMK-1/p38 MAPK activation and drive the expression of immune genes.

To extend our analysis, we compared our dataset to a previous study that identified genes differentially expressed in animals acclimated to 21% and 7% O_2_^29^. Most of the neuropeptides regulated by IL-17 were not regulated by O_2_ experience^29^ (Supplementary Table 3), suggesting IL-17 elicits transcriptional changes not explained by altered activity of the O_2_-sensing circuit. These data suggested that IL-17 signaling may directly or indirectly alter many features of *Ce* behavior and global physiology.

### Neural IL-17–MALT-1 signaling suppresses immunity and reduces lifespan

To explore immune functions of IL-17 we measured survival on *Pseudomonas aeruginosa*, a bacterial pathogen that colonizes the intestine of *C. elegans*^44^. We carried out these experiments in animals having the N2 version of the *npr-1* neuropeptide receptor, *npr-1 215V*, which inhibits aggregation behavior and escape from 21% O_2_^32, 45^; this ensures differences in hyperoxia avoidance do not contribute to altered pathogen resistance.

Animals lacking *malt-1,* or harboring the *malt-1* protease-dead allele *malt-1(C374A),* were resistant to PA14 infection compared to N2 controls (Fig 6c and d). PA14 resistance in *malt-1(null)* mutants was rescued by pan-neuronal expression of *malt-1* cDNA (Fig. 6c), suggesting that MALT-1 acts in the nervous system to regulate the immune response. Like *malt-1* mutants, *ilcr-1* and *nfki-1* mutants survived signficantly longer on PA14 than controls (Fig. 6e, f).

*Ce* innate immune responses to *P. aeruginosa* are regulated by both insulin/insulin-like growth factor signaling (IIS) and the p38 MAPK pathway^46^. The p38 MAPK pathway functions downstream of TIR-1 (Toll/Interleukin-1 Receptor domain protein), the *C. elegans* ortholog of SARM1 (Sterile alpha and TIR motif containing protein)^24, 25^. MALT-1 strongly and specifically co-IP’d TIR-1 (Supplementary Table 1d). TIR-1 regulates expression of anti-microbial peptides, including the ShK-like toxin T24B8.5 in the intestine^47, 48^. Like *tir-1* mutants^48^, *malt-1* mutants showed reduced T24B8.5 expression (Supplementary Fig. 7a). This reduction was rescued by intestine-specific or nervous system-specific expression of *malt-1* (Supplementary Fig. 7A).

TIR-1 regulates T24B8.5 expression by promoting phosphorylation of the p38 MAPK PMK-1^25^. Like *tir-1* mutants, *malt-1* mutants showed reduced PMK-1 phosphorylation (Supplementary Fig. 7b). Therefore, mutants defective in IL-17/MALT-1 signaling exhibit PA14 resistance despite having apparently reduced PMK-1 activity. PA14 resistance was, however, greatly reduced in *malt-1; tir-1* double mutants compared to *malt-1* (Fig. 6g), indicating that TIR-1/SARM can still promote PA14 resistance in *malt-1* mutants. Together, these data indicate that while overall IL-17 signaling negatively regulates the *Ce* immune response to PA14, this effect may reflect the net outcome of opposing influences.

To determine whether IL-17 mutants alter lifespan, as well as pathogen resistance, we measured survival on the standard laboratory food source of *C. elegans*, *E. coli* OP50. *ilc-17.1* and *malt-1* mutants lived significantly longer than N2 controls (Fig. 6h, i). Expression of *ilc-17.1* cDNA from its endogenous promoter not only rescued the phenotype of the null mutant, but significantly reduced lifespan compared to non-transgenic N2 controls (Fig. 6h). We could also rescue the extended lifespan of *malt-1* mutants by pan-neuronal expression of *malt-1* gDNA (Fig. 6i), suggesting that IL-17 signaling acts in the nervous system to regulate longevity.

### IL-17 signaling modulates associative learning

The widespread expression of *malt-1* and IL-17 pathways genes in the nervous system made us speculate that IL-17 signaling shapes other behaviors. We previously showed that mutants in *ilc-17.1*, *pik-1* and *nfki-1* retained normal basal responses to salt and odors^17^. We wondered, however, if IL-17 signals might alter how experience shapes sensory responses. To test this hypothesis, we used a salt chemotaxis paradigm, in which associating an environment high in NaCl with food withdrawal, leads to decreased salt attraction when animals are subsequently tested in a chemotaxis assay^49^. All *Ce* IL-17 mutants tested showed stronger attraction to salt than N2 controls shortly after conditioning (Fig. 6j). We could rescue the *ilcr-1* phenotype by selectively expressing cDNA encoding the ILCR-1 receptor in the ASE salt-sensing neurons (*flp-6p*), but not in the RMG interneurons (*flp-5p*) or the O_2_ sensors (*gcy-32p*) (Fig. 6k). Thus, the IL-17 pathway promotes salt chemotaxis learning, and IL-17 signaling in the nervous system is not restricted to the O_2_-sensing circuit.

In summary, our data suggest that IL-17 signals through a MALT-1 signalosome to modify neural properties and remodel the behavior and physiology of *C. elegans*.

## Discussion

Our data show that MALT1 modulates neural circuit function in *C. elegans,* by acting as a protease and a scaffold. MALT-1 forms an ACTL-1-IRAK-MALT-1 signaling complex that mediates IL-17 signaling. The high molecular weight of this complex in *Ce* extracts suggests it may form a structure related to the MYD88-IRAK4-IRAK2 Myddosome^39, 40^ and CARMA1-BCL10-MALT1 CBM signalosome (Qiao et al., 2013). MALT-1 directly binds ACTL-1 *in vitro*, and yeast two hybrid data suggest ACTL-1 directly binds *Ce* IRAK^50^. MALT-1 also interacts directly with NFKI-1, a homolog of mammalian IκBζ/IκBNS, and can signal through both NFKI-1-dependent and independent mechanisms to alter neuron function and change behavior. The ACTL-1-IRAK- MALT-1-NFKI-1 pathway is present in most neurons of the *C. elegans* nervous system, and appears to mediate a major neuromodulatory axis that impacts multiple phenotypes.

IL-17Rs are found throughout metazoa^51^. We speculate that the ACTL-1-IRAK-MALT-1 complex we have identified is the original and primary mechanism by which IL-17Rs signal in non-amniote animals, from cnidarians to cephalochordates. In amniotes, ACT1 orthologs have lost a death domain (DD) that is present in ACT1 orthologs from other lineages^51^. DD mediate homotypic interactions in large immune complexes such as the Myddosome^52^, and are present in both MALT1 and IRAKs. The DD – SEFIR domain architecture of ACT1 present in non-amniotes resembles the DD – TIR domain structure of MYD88, since TIR and SEFIR domains are related^53^. Interestingly, proximity labelling studies find MALT1 associates with MYD88 in DLBCL cells^54^. Recent studies have also speculated that mammalian MALT1 is recruited to IL-17 signaling complexes^55, 56^. Direct evidence for this is lacking, but if correct, this would mirror our results in *C. elegans*.

One area of future investigation will involve studying how MALT-1 alters neural function. Although we find that MALT-1 protease activity is essential, we have not identified its neural substrate(s). Substrates of mammalian MALT1 with orthologs in *C. elegans* include the RNA binding proteins roquin-1/2 (RLE-1), which target RNAs for degradation, the endoribonuclease regnase-1 (REGE-1), and the CYLD (CYLD-1) deubiquitinase^57–59^. We did not detect specific behavioral defects in mutants for these genes (unpublished data). However, our IP/MS data indicate that besides binding NFKI-1, MALT-1 interacts with multiple RNA binding proteins, including splicing and polyadenylation factors. A plausible hypothesis is that in the nervous system, like in the mammalian immune system, MALT-1 can regulate gene expression at the RNA level. MALT-1 also binds the *Ce* ortholog of SARM1, called TIR-1. TIR-1 regulates *Ce* gustatory and olfactory plasticity^27^, and proteostasis^60^, by modulating MAPK pathways, making it a plausible target for regulation by MALT-1.

The closest mammalian homolog of NFKI-1, IκBζ, is a nuclear-localized protein that acts as a transcriptional regulator, and is rapidly induced by inflammatory stimuli, including IL-17. IκBζ is thought to mediate its effects on gene expression primarily by regulating chromatin structure, although how it is recruited to target genes is not completely understood since it lacks a DNA binding domain^61, 62^. Our IP/MS data find NFKI-1 physically interacts with the CREB binding protein (CREBBP), *cbp-1*, which is a histone acetyltransferase, consistent with NFKI-1 acting to modify chromatin structure. While our RNA-seq studies may highlight some genes regulated by NFKI-1, most of the gene expression changes we report likely reflect secondary consequences of signalling defects. Interestingly, the small number of specific MALT-1 interactors we identify by IP/MS includes the nuclear hormone receptors NHR-49 and NHR-71. NHR-49 is the *Ce* ortholog of hepatocyte nuclear factor 4A/G^63^.

Our biochemical and genetic analyses of IL-17 signaling in the *Ce* model have identified functional roles and biochemical interactions previously undescribed in mammals. An outstanding challenge is to examine which of these are conserved in mammals. To name a few examples, does MALT1 play a role in modulating mammalian neural function, given that neurons can express receptors with an ITAM (immunoreceptor tyrosine-based activation motif)^18–20^, as well as GPCRs, that in immune cells signal through MALT1? Does MALT1 contribute to known neuronal responses to IL-17? Does mammalian MALT1 physically interact with IκBζ, IκBNS, or SARM1? In summary, we have used an invertebrate model system, with the relative simplicity this offers, and its advantages for genetics, biochemistry and single neuron analysis, to probe how key immune molecules signal in neurons to alter circuit function.

## Acknowledgements

We thank the *Caenorhabditis* Genetics Center (funded by National Institutes of Health Infrastructure Program P40 OD010440) and the Japanese knockout consortium for strains, the Cambridge Research Institute Genomics Core for whole genome sequencing, Adeline Colussi and Harvey McMahon for reagents, and de Bono lab members for comments on the manuscript. This work was supported by the Medical Research Council UK, the European Research Council (Advanced Grant 269058 to M.d.B), and Wellcome (209504/Z/17/Z Investigator Award to M.d.B.).

## Author contributions

S.F., C.C., M.A., S.B. and M.d.B. conceived experiments; S.F., C.C., M.A., and S.B. performed experiments; A.C. and G.N. performed sequence analysis; S-Y C-P and F.B performed mass spectrometry analysis; S.F. and M.d.B. wrote the manuscript.

## Supplementary Figure Legends

**Supplementary Figure 1.**
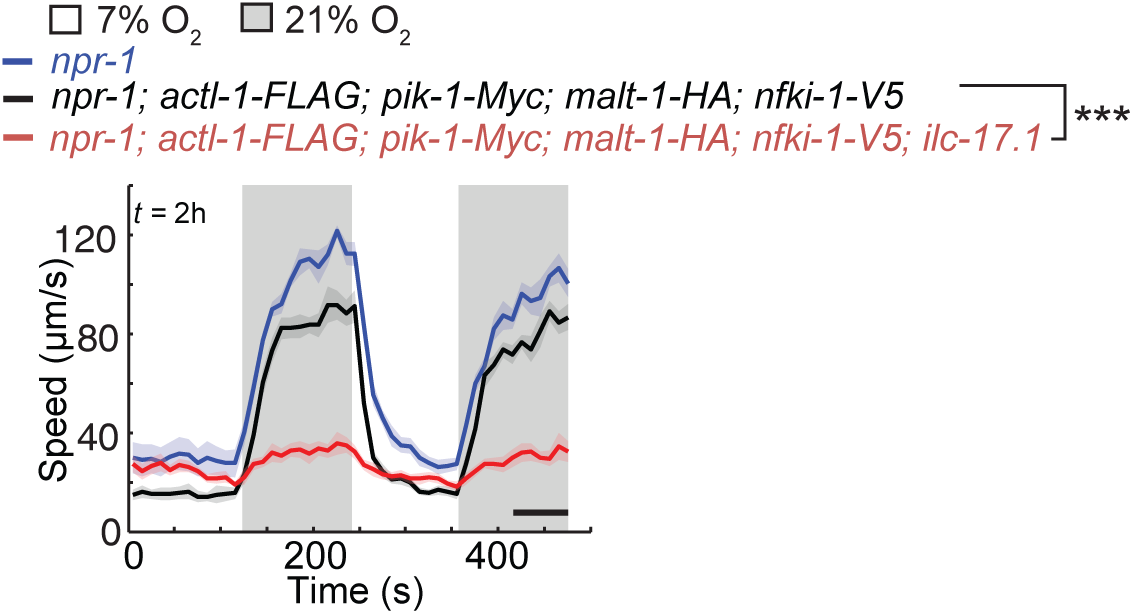
Related to Figure 1. Knocking in epitope tags into endogenous *actl-1*, *pik-1*, *malt-1*, and *nfki-1* does not disrupt their function. A strain co-expressing *actl1-1-HA*, *pik-1-Myc*, *malt-1-HA* and *nfki-1-V5,* in an *npr-1* background, exhibits similar arousal to *npr-1* controls in response to 21% O_2_. Arousal is abolished when these knock-in alleles are crossed into an *ilc-17.1(tm5218)* background. ***, P < 0.001, Mann-Whitney *U* test, n ≥ 81 animals.

**Supplementary Figure 2.**
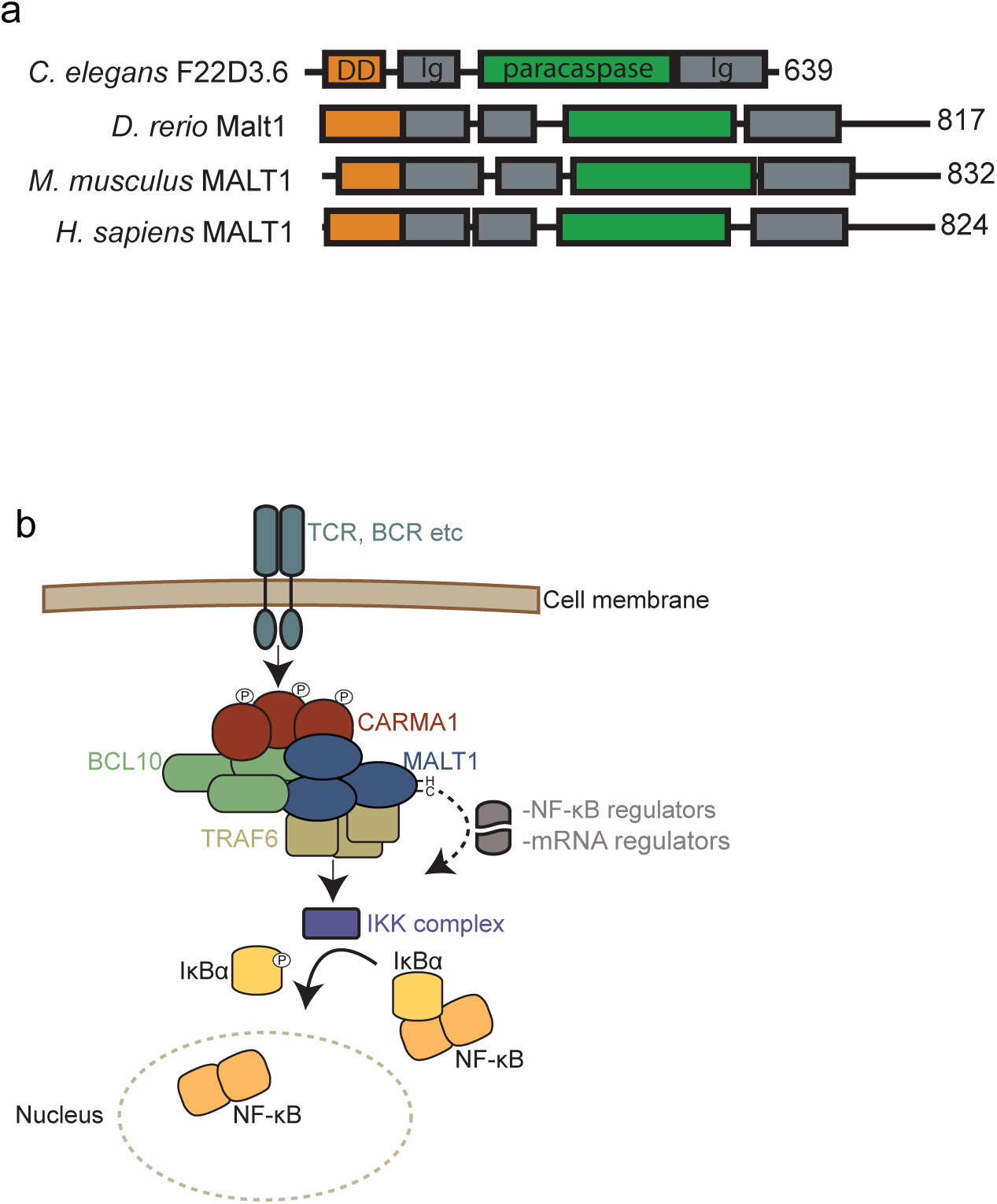
Related to Figure 1. MALT1. **a** MALT1 paracaspase domain organization. DD,death domain; Ig, Immunoglobulin-like fold. **b** Schematic of MALT1 signaling in mammalian B and T cells. BCR/TCR stimulation induces formation of the CARMA1-BCL10-MALT1 (CBM) signalosome via protein kinase C β (PKCβ)/PKCθ-mediated phosphorylation of CARMA1. MALT1 recruits TRAF6, whose ubiquitin ligase activity leads to the recruitment and activation of the IKK complex. This culminates in IKK-mediated phosporylation of IκB*α*, triggering its degradation and releasing NF-κB for nuclear translocation. Additionally, MALT1 promotes lymphocyte activation by cleaving negative regulators of NF-κB, and factors that regulate mRNA stability. Blastp suggests nematode genomes do not harbor orthologs of BCL10 or CARMA.

**Supplementary Figure 3.**
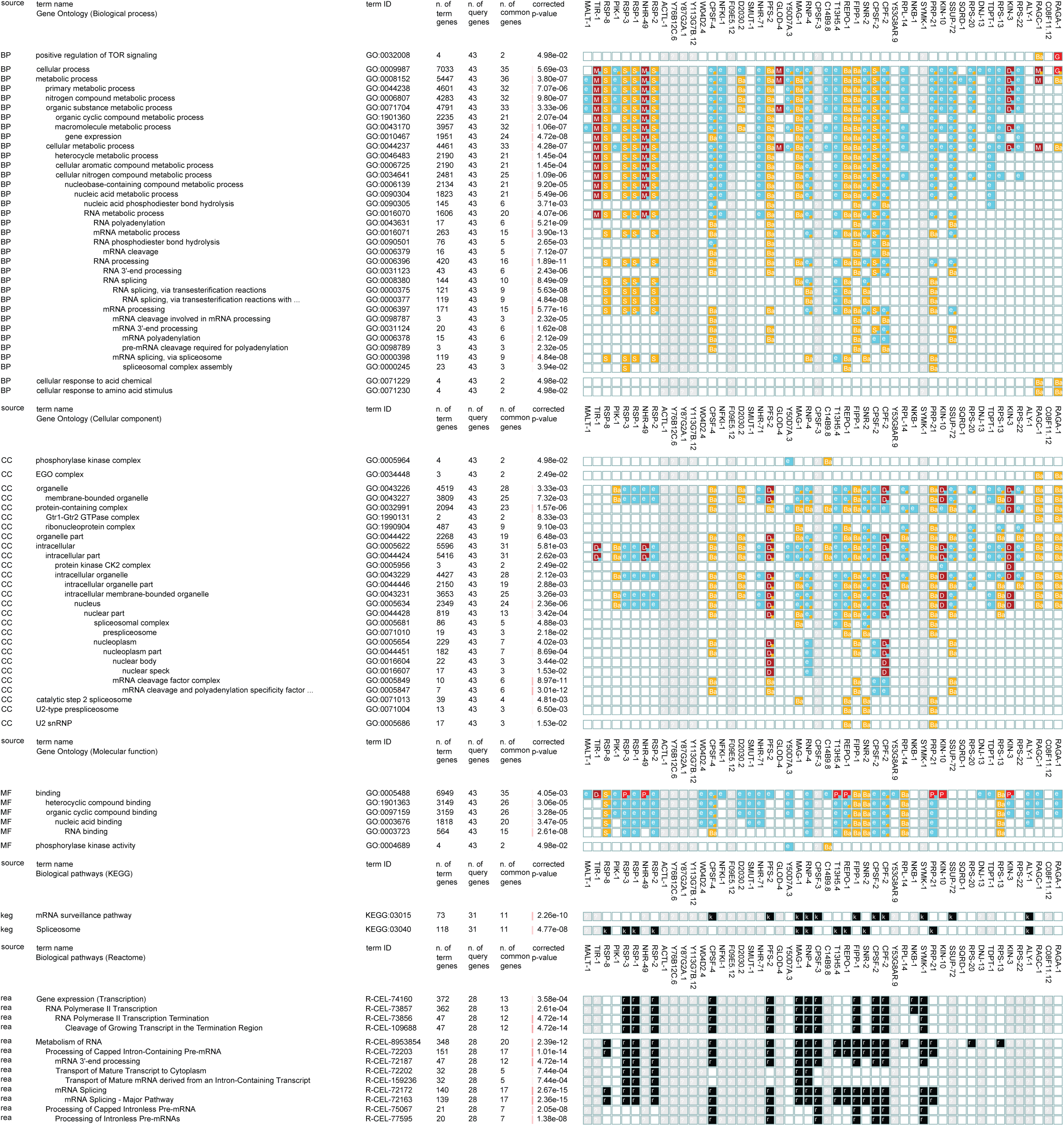
Related to Figure 1. The MALT-1-interacting proteome is enriched in factors involved in RNA metabolism. Gene ontology and biological pathway enrichment analysis of the 50 factors that specifically co-immunoprecipitate with MALT-1 (Supplementary Table 1), performed with g:Profiler^65^.

**Supplementary Figure 4.**
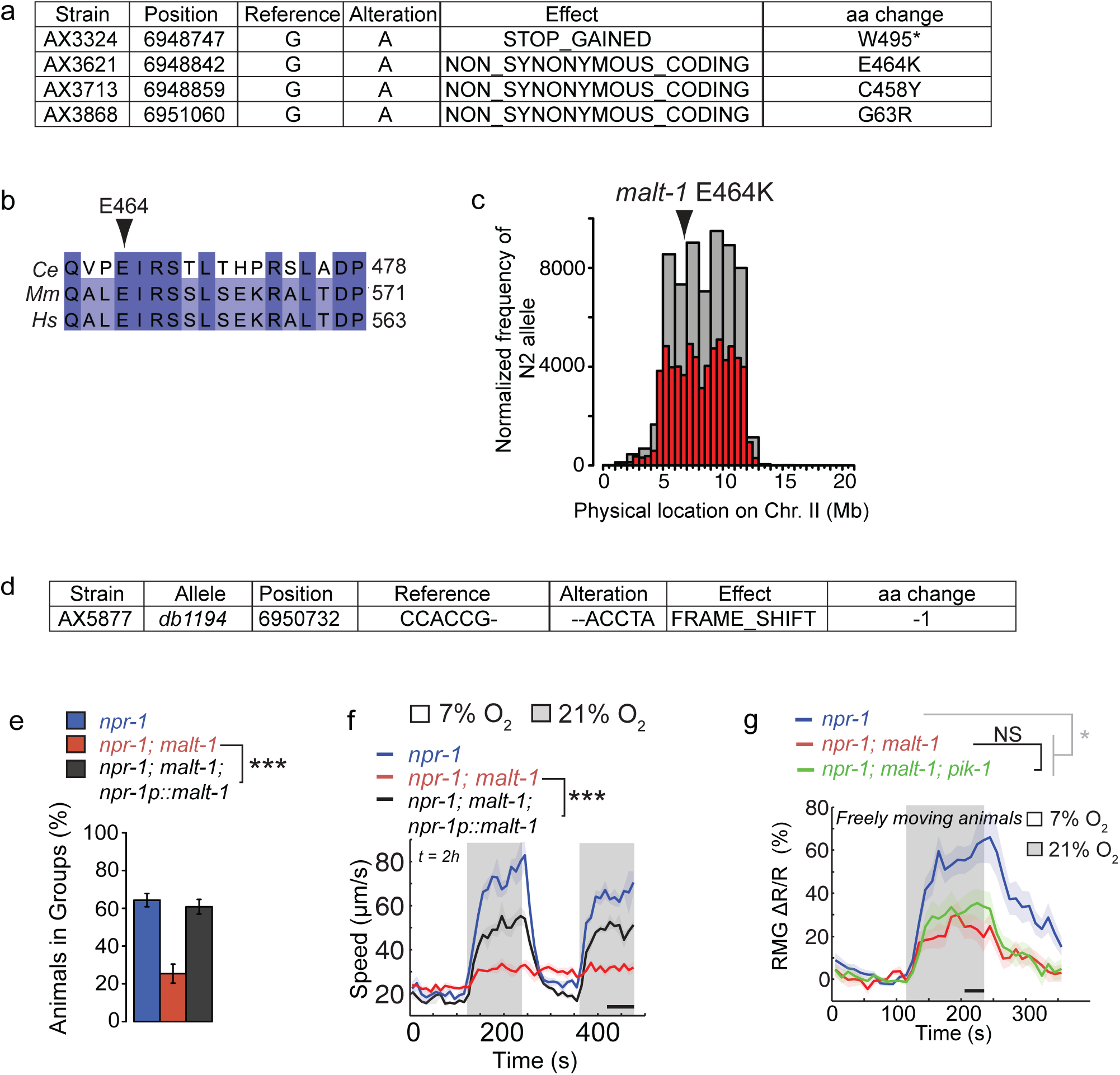
Related to Figure 1. MALT-1 promotes escape from 21 O_2_. **a** Mutations in *malt-1* isolated in a forward genetic screen for loss of aggregation behavior. **b** The MALT-1 E464 residue mutated in AX3621 is conserved from *Ce* to human, and corresponds to E549 in human. *Mm* = *Mus musculus, Hs* = *Homo sapiens*. **c** The aggregation defect of AX3621 maps close to *malt-1*. Mapping used CloudMap (see Methods) which measures the relative levels of SNPs derived from two genetic backgrounds in aggregation-defective recombinants, N2 single nucleotide polymorphisms (SNPs) are derived from AX3621; Hawaiian SNPs are derived from the AX288 (*lon-2 npr-1*) Hawaiian background strain. Aggregation-defective recombinants show an enrichment of N2 Bristol SNPs on chromosome II, flanking the physical location of the *malt-1* mutation at 6.9 Mb. **d** The *malt-1(db1194)* allele, generated by CRISPR/Cas9 cleavage at the *malt-1* locus. **e** and **f** The aggregation phenotype (**e**) and arousal defect in 21% O_2_ (**f**) of *malt-1(db1194)* mutants is rescued by expressing *malt-1* cDNA from the *npr-1* promoter, which drives expression in a broad subset of neurons including URX and RMG^28, 45^. n ≥ 44 animals, ***, P < 0.001, Mann-Whitney *U* test. **g** *malt-1* RMG Ca^2+^ transients (reported by YC2.60) in freely moving animals are not further reduced by loss-of-function mutations in *pik-1* (n ≥ 14 animals). *, P < 0.05, Mann-Whitney *U* test.

**Supplementary Figure 5.**
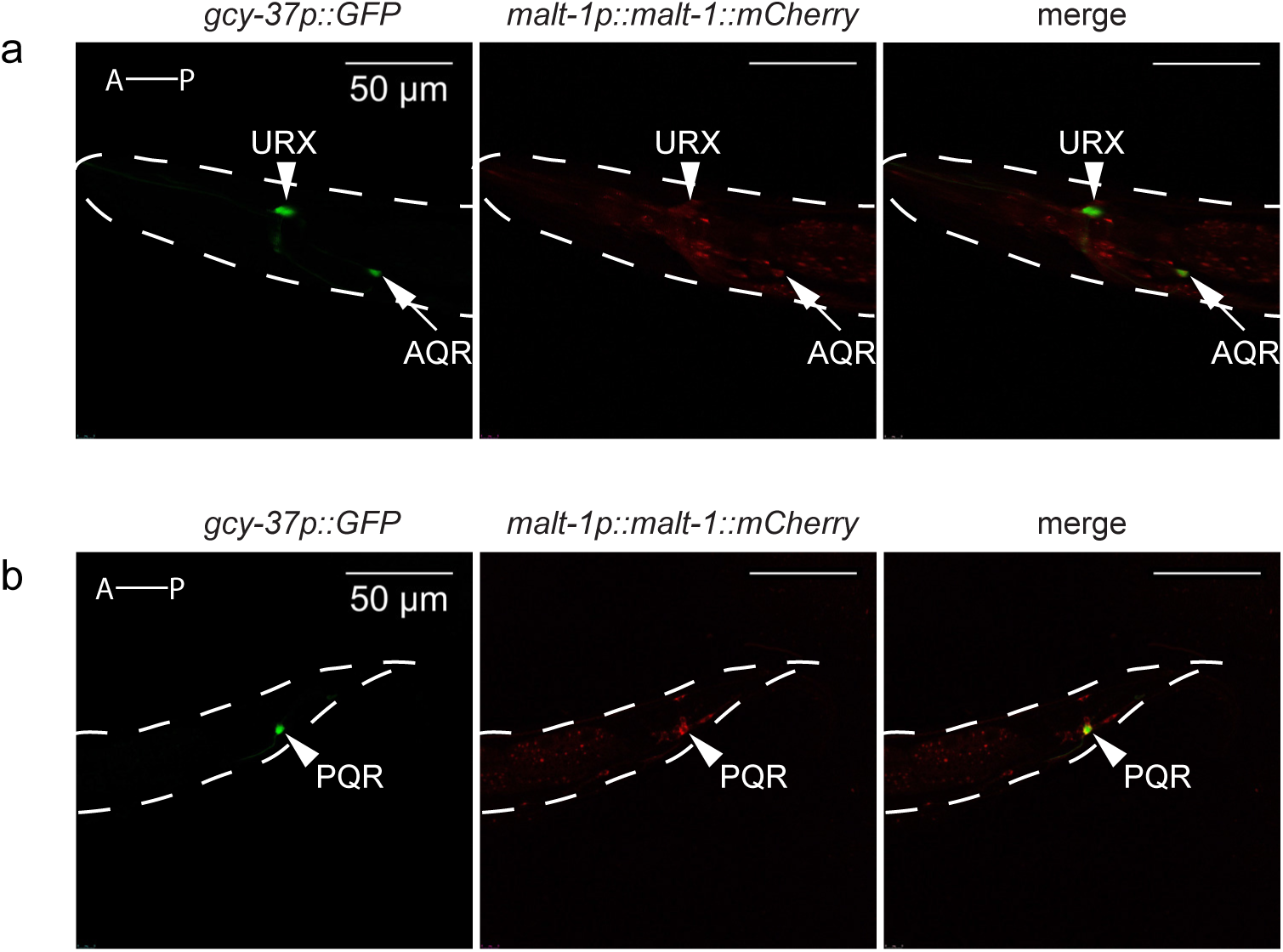
Related to Figure 2. MALT-1 is expressed in O_2_-sensing neurons. A MALT-1::mCherry translational fusion, expressed from its endogenous promoter (4kb), is expressed in URX and AQR in the head (**a**) and PQR in the tail (**b**), which are labelled with a *gcy-37::gfp* reporter.

**Supplementary Figure 6.**
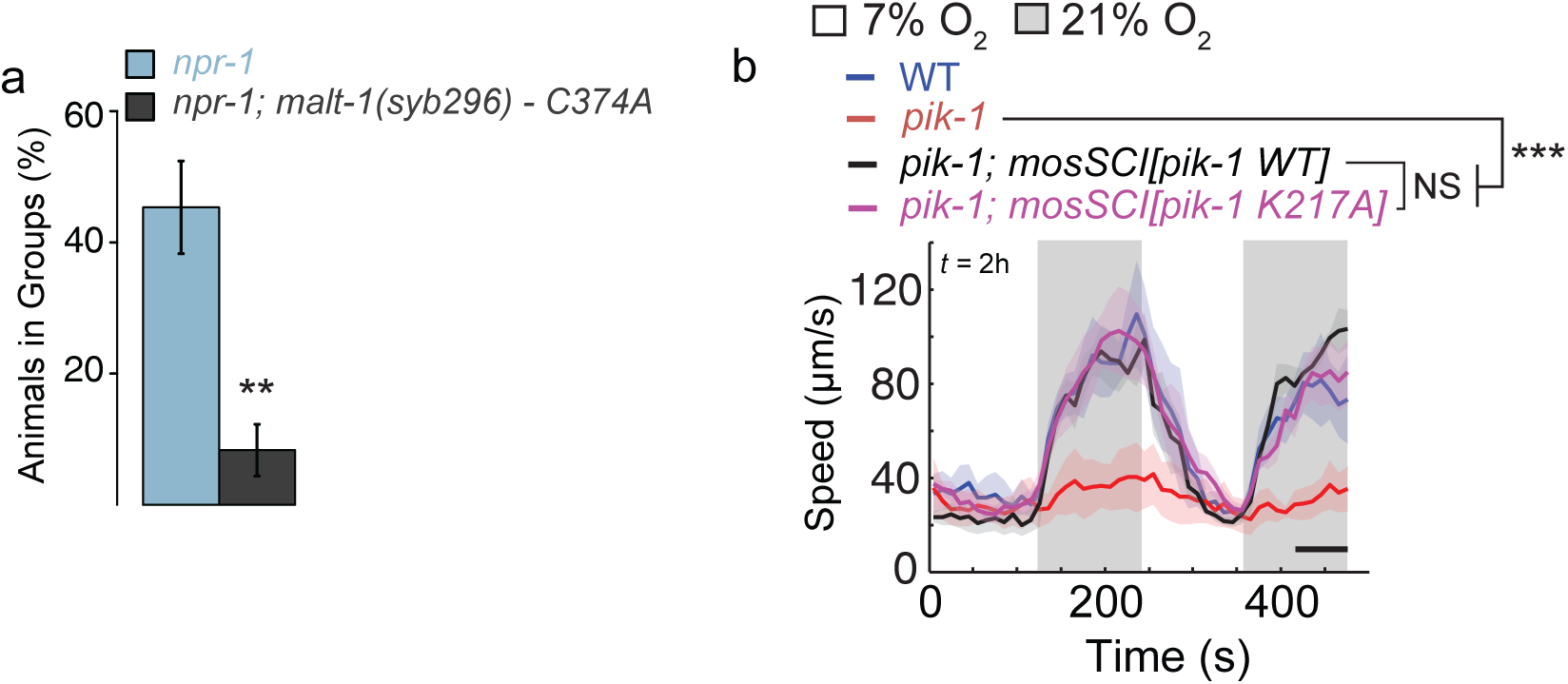
Related to Figure 4. Targeted disruption of the protease activity of MALT-1 and kinase activity of PIK-1/IRAK. **a** *malt-1(syb296)* mutants expressing catalytically inactive MALT-1 C374A exhibit strong aggregation defects compared to *npr-1* animals. **b** PIK-1 ATP-binding may not be required for avoidance of 21% O_2_. Single copy transgenes (MosSCI) expressing *pik-1* WT and *pik-1 K217A* cDNA rescue the O_2_-response defect of *pik-1(tm2167)* mutants equally well. K217 corresponds to the lysine residue coordinating ATP binding in kinase active sites (n ≥ 40 animals). ***, P < 0.001, Mann-Whitney *U* test.

**Supplementary Figure 7.**
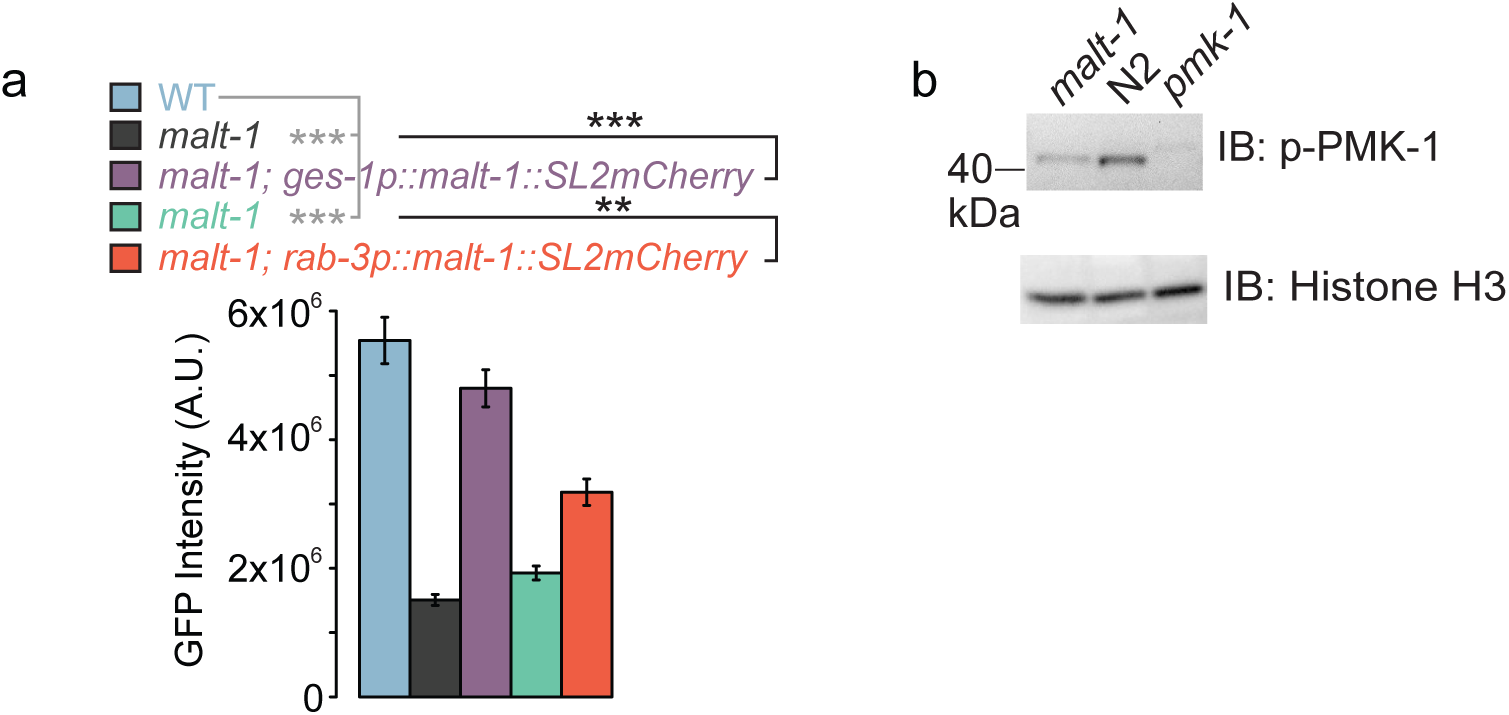
Related to Figure 6. MALT-1 promotes p38 MAPK signaling. **a** Expression from a reporter gene, *T24B8.5p::GFP,* stimulated by PMK-1 activity. GFP expression is decreased in *malt-1(db1194)* mutants compared to N2, and restored by tissue-specific expression of WT *malt-1* cDNA in the intestine (*ges-1* promoter) or the nervous system (*rab-3* promoter). n ≥ 24 animals, **, P < 0.01, ***, P < 0.001, ANOVA with Tukey’s post hoc HSD. **b** Immunoblot analysis comparing levels of phosphorylated PMK-1 in *malt-1* and *ilc-17.1* mutants to N2 controls. Histone H3 is analyzed in parallel as a loading control.

## Supplementary Tables

**Supplementary Table 1** Related to Figure 1. LC-MS/MS analysis of proteins co-immunoprecipitating with IL-17 signaling components. Shown are total spectral counts for all proteins that specifically co-IP with ACTL-1(**a**) PIK-1(**b**) NFKI-1(**c**) and MALT-1(**d**).

**Supplementary Table 2** Related to Figure 6. Gene-expression profiles of *ilc-17.1(tm5218)*; *npr-1, malt-1(db1194)*; *npr-1,* and *nfki-1(db1197); npr-1* mutants compared to *npr-1* controls.

**a** and **b** All genes significantly downregulated (**a**) and upregulated (**b**) in all three conditions compared to control. FPKM values, fold-change and statistics for each gene are shown for the *npr-1* vs *npr-1; ilc-17.1(tm5218)* comparison.

**c** and **d** Significantly enriched GO (**c**) and KEGG (**d**) terms in genes whose expression is significantly altered in all three mutants compared to control, with a log_2_ fold-change cut-off of 0.25.

**Supplementary Table 3** Related to Figure 6. Comparison of neuropeptide gene expression in mutants defective in response to 21% O_2_^29^, and mutants defective in IL-17 signaling.

**Supplementary Table 4.**
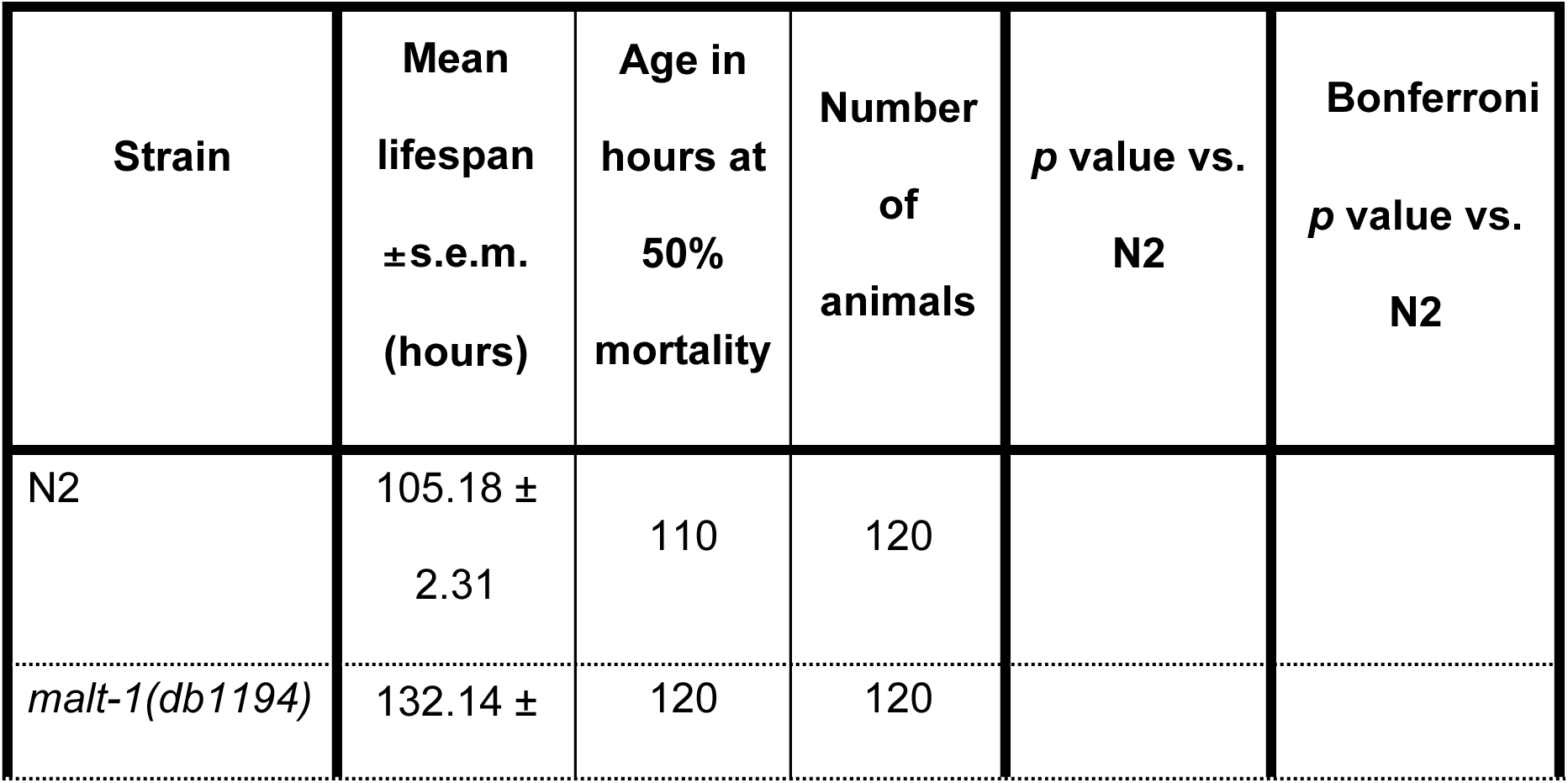
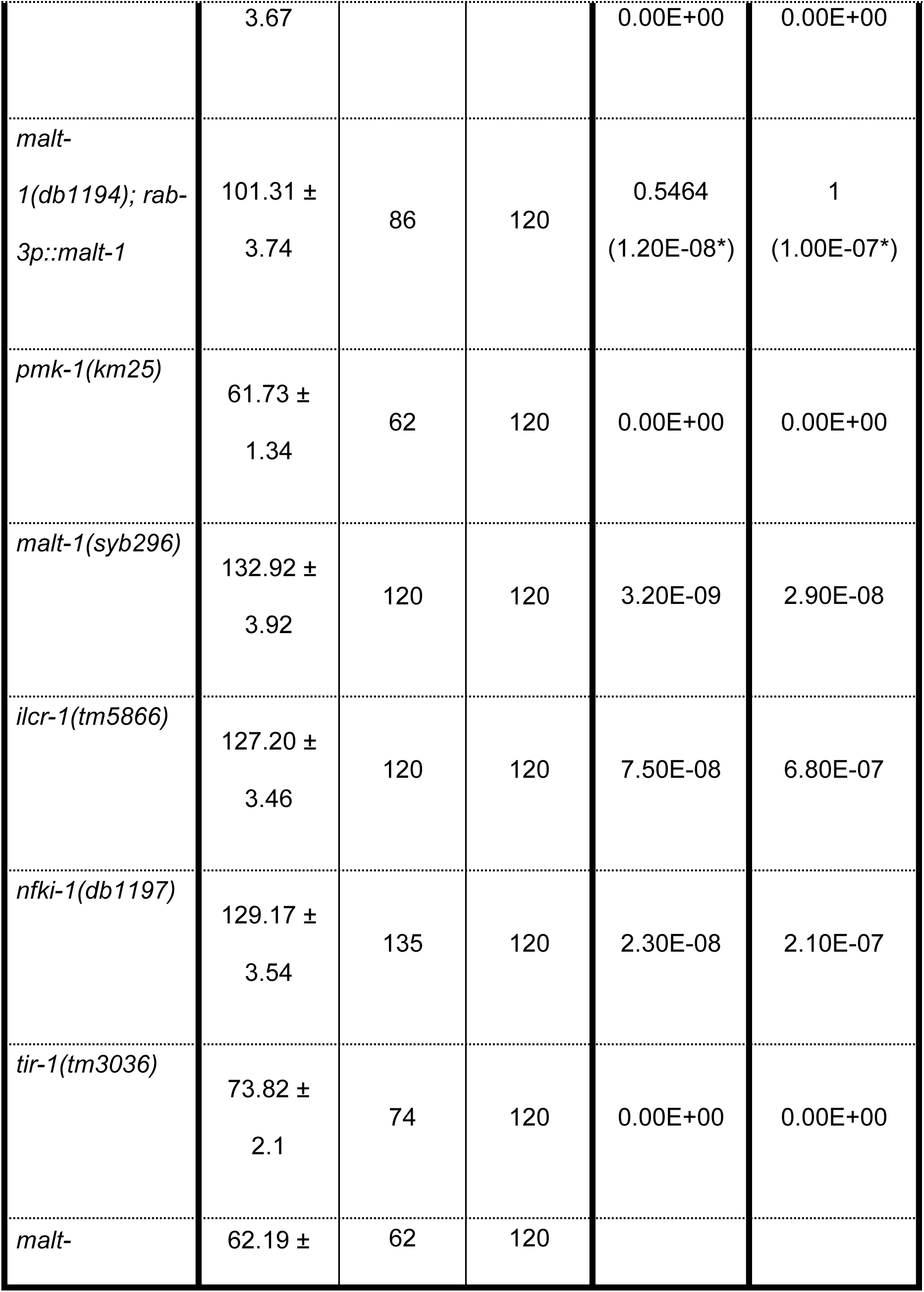
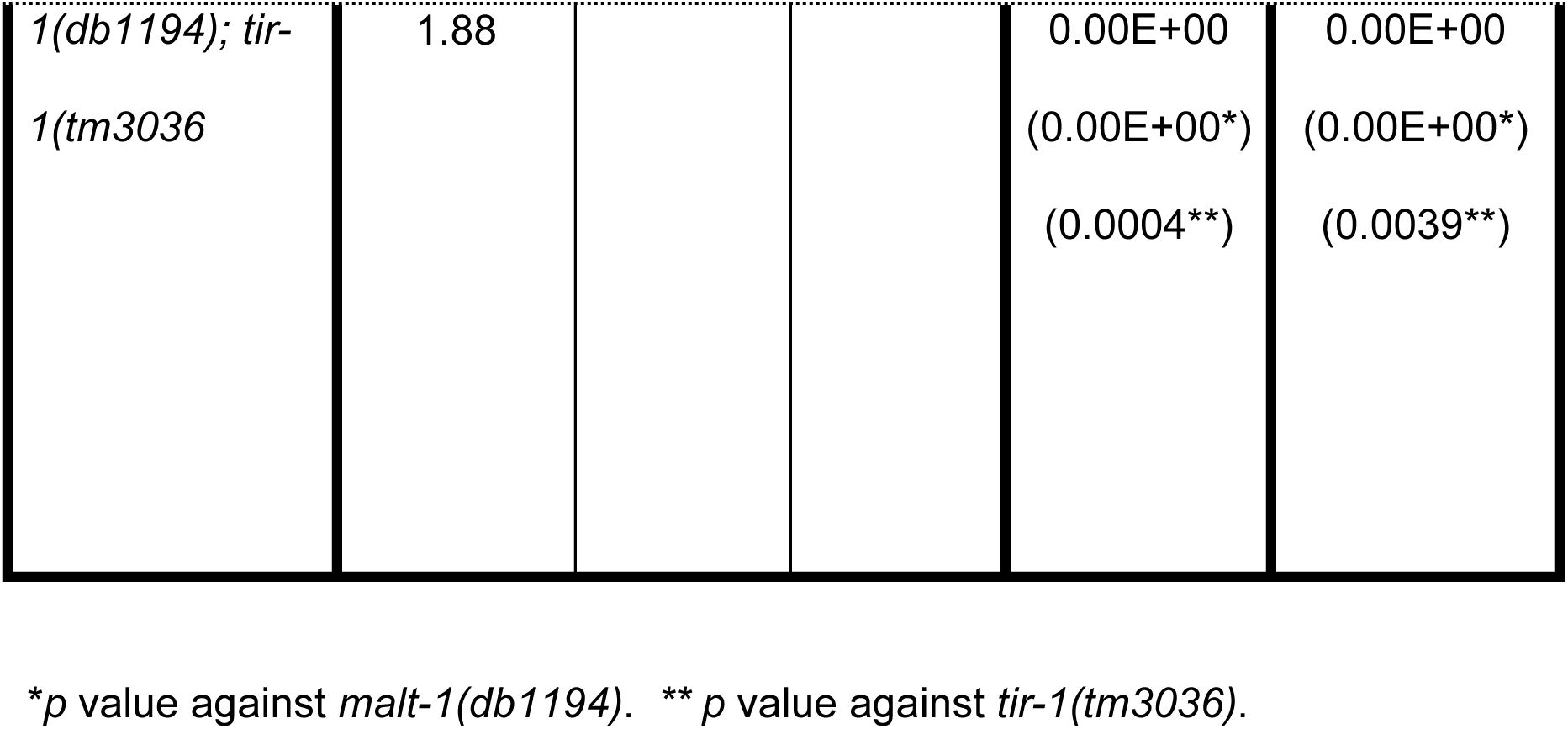
Related to Fig. 6. PA14 Survival data analysis of IL-17 pathway mutants.

**Supplementary Table 5.** Related to Fig. 6. Lifespan of IL-17 pathway mutants on OP50.

| Strain | Mean lifespan<br>±s.e.m.<br>(days) | Age in days at 50% mortality | Number of animals | *p* value vs. N2 | Bonferroni *p* value vs. N2 |
| --- | --- | --- | --- | --- | --- |
| N2 | 17.32 ± 0.37 | 17 | 180 |  |  |
| *malt-1(db1194)* | 20.84 ± 0.58 | 21 | 180 | 7.40E-08 | 2.90E-07 |
| *malt-1(db1194); rab-3p::malt-1* | 18.78± 0.46 | 19 | 180 | 0.0026<br>(0.0029*) | 0.0105<br>(0.0115*) |
| *ilc-17.1*<br>(*tm5218*) | 19.00 ±<br>0.45 | 19 | 180 | 0.0024 | 0.0097 |
| *ilc-17.1*<br>(*tm5218*); *ilc-17.1p::ilc-17.1* | 13.84 ±<br>0.31 | 15 | 180 | 0.00E+00<br>(0.00E+00**) | 0.00E+00<br>(0.00E+00**) |

\**p* value against *malt-1(db1194)*. \*\**p* value against *ilc-17.1(tm5218)*.

**Supplementary Table 6.**
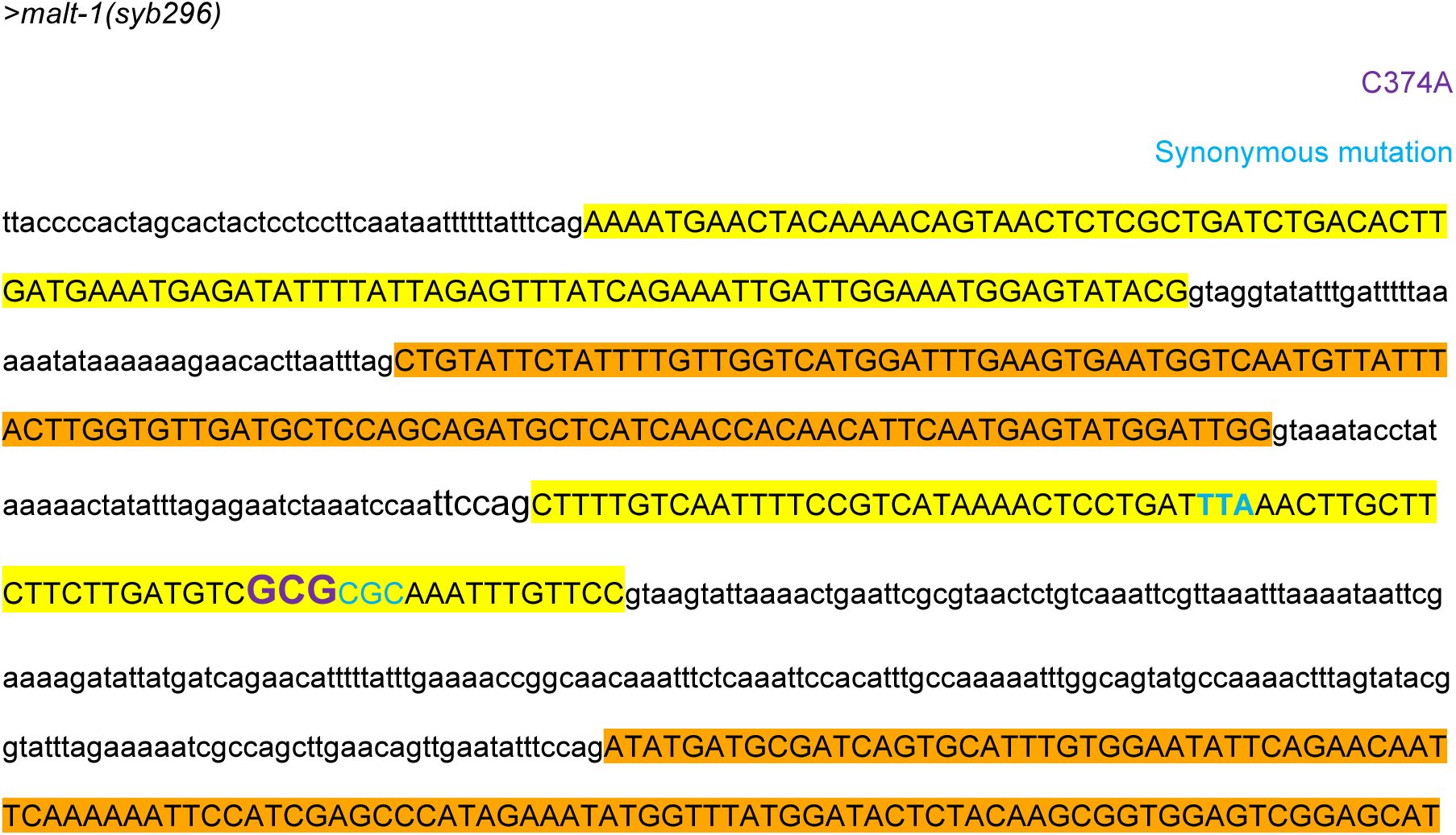
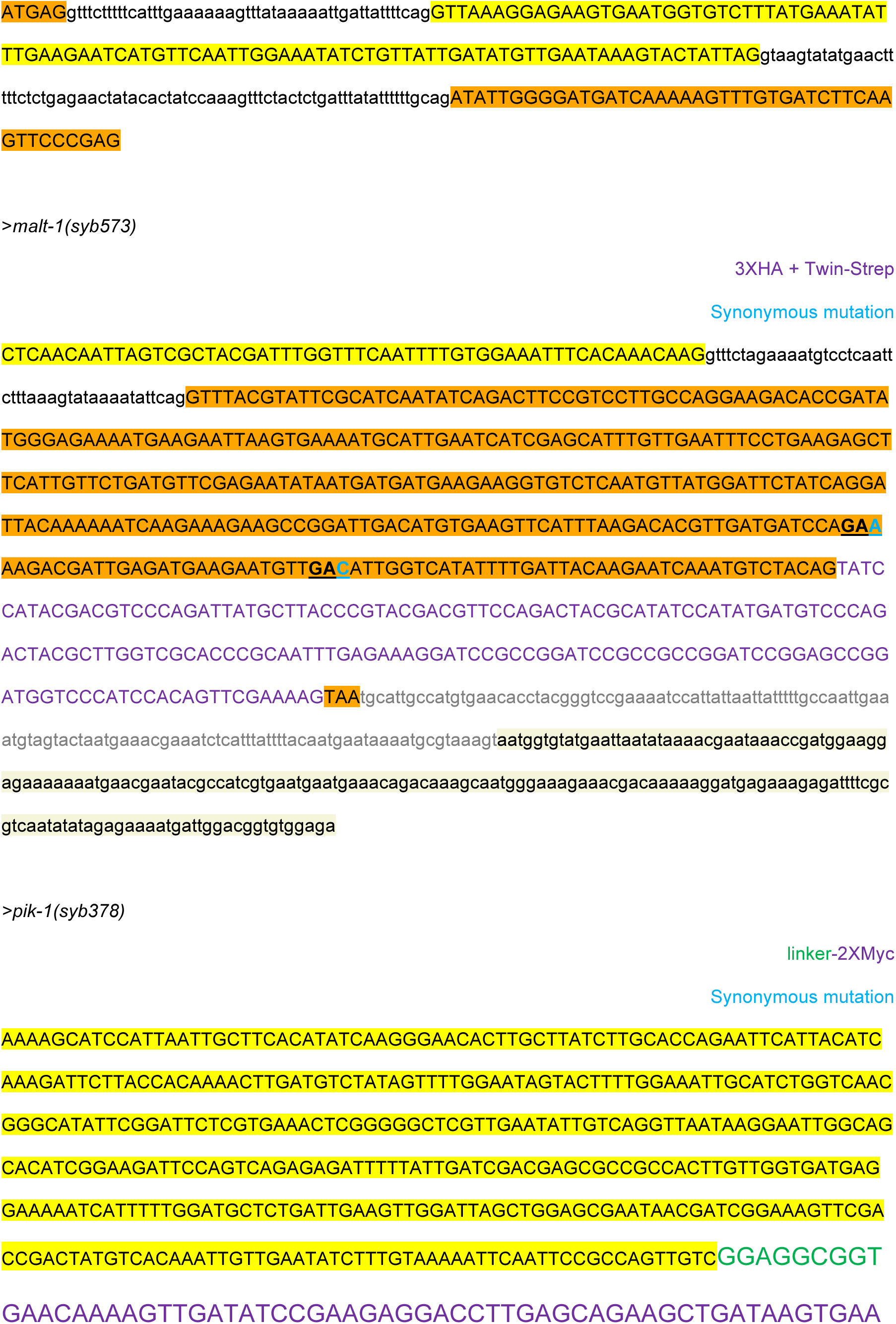
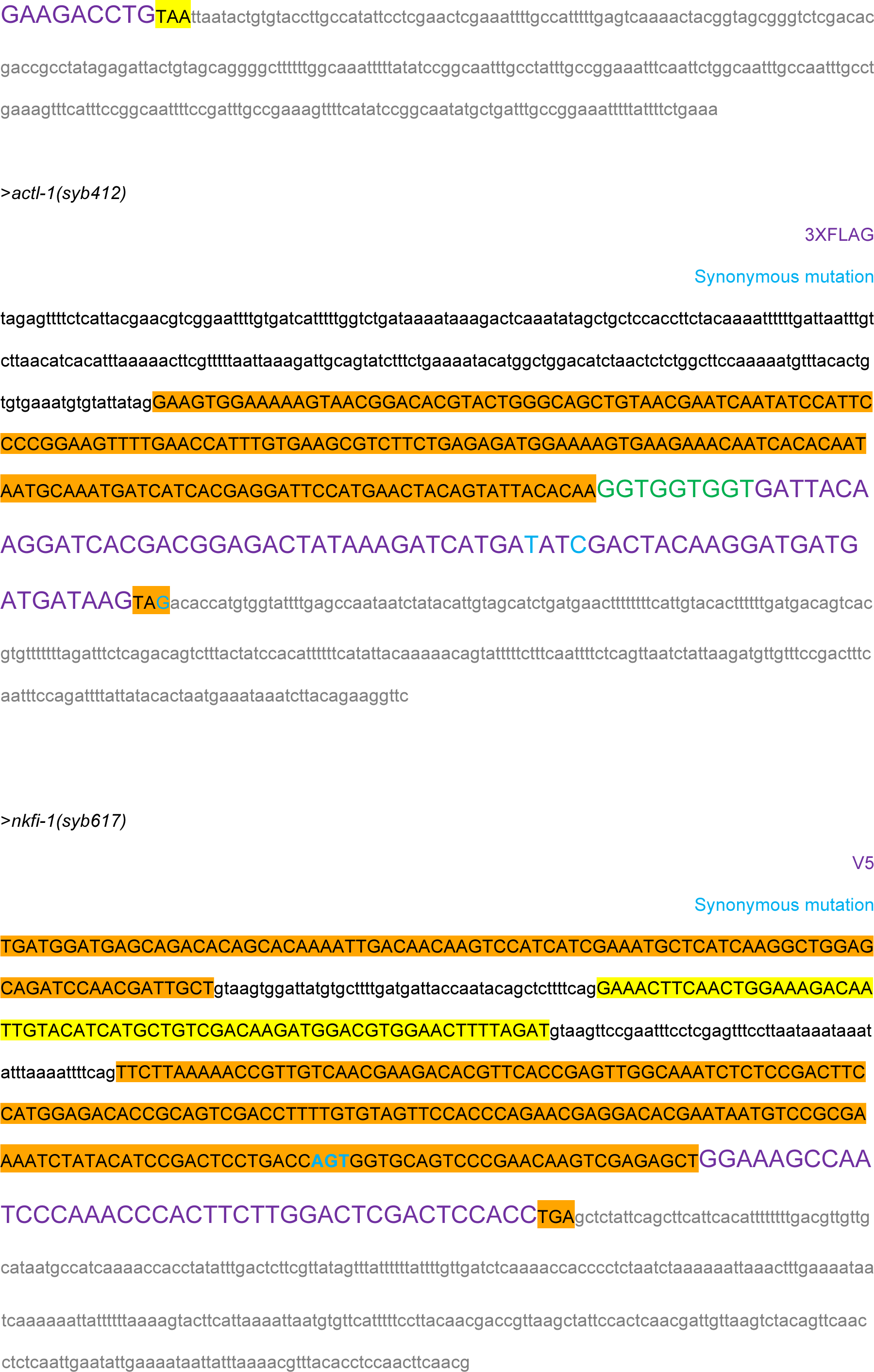
Knockin alleles used in this study.

**Supplementary Table 7.**
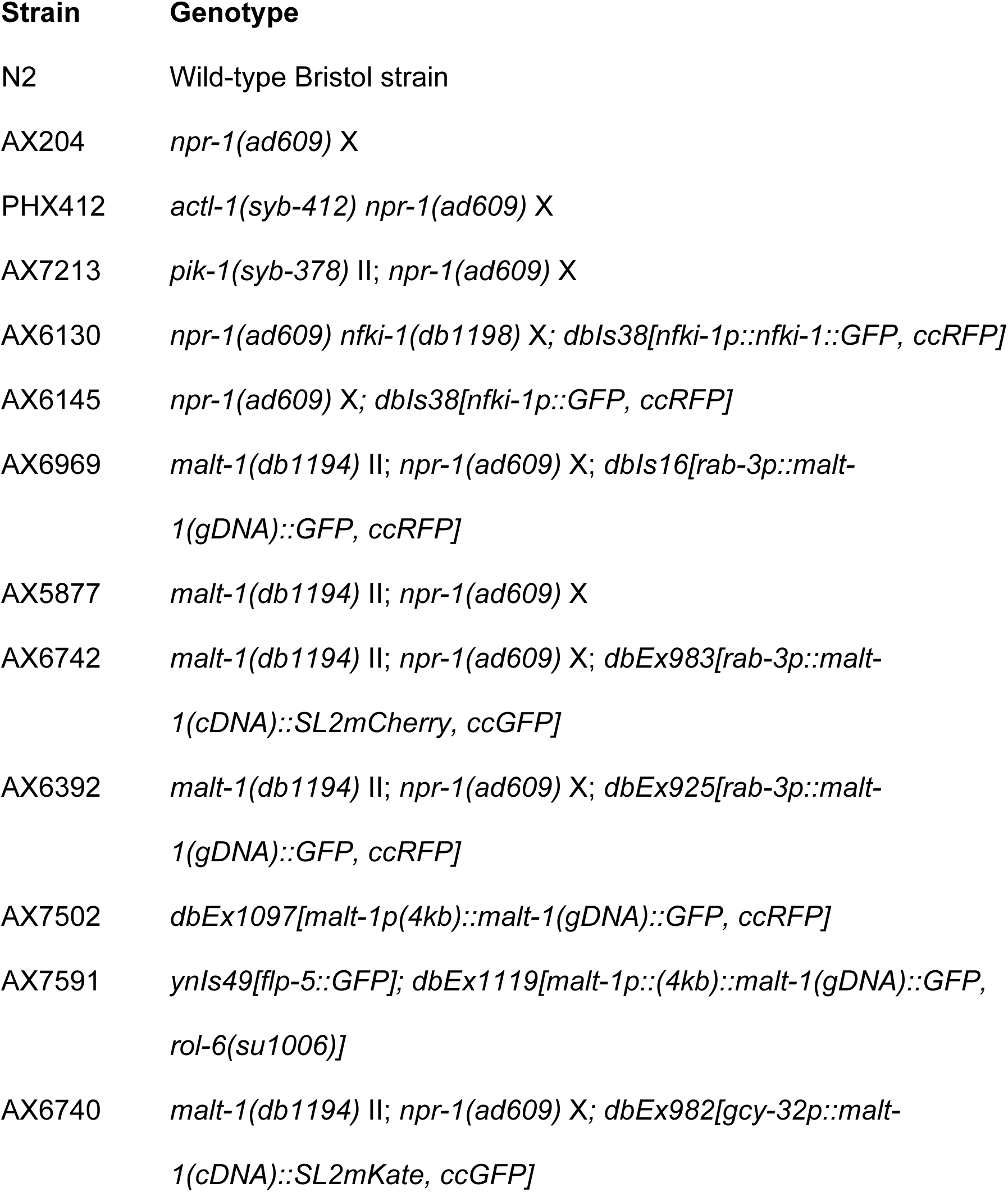
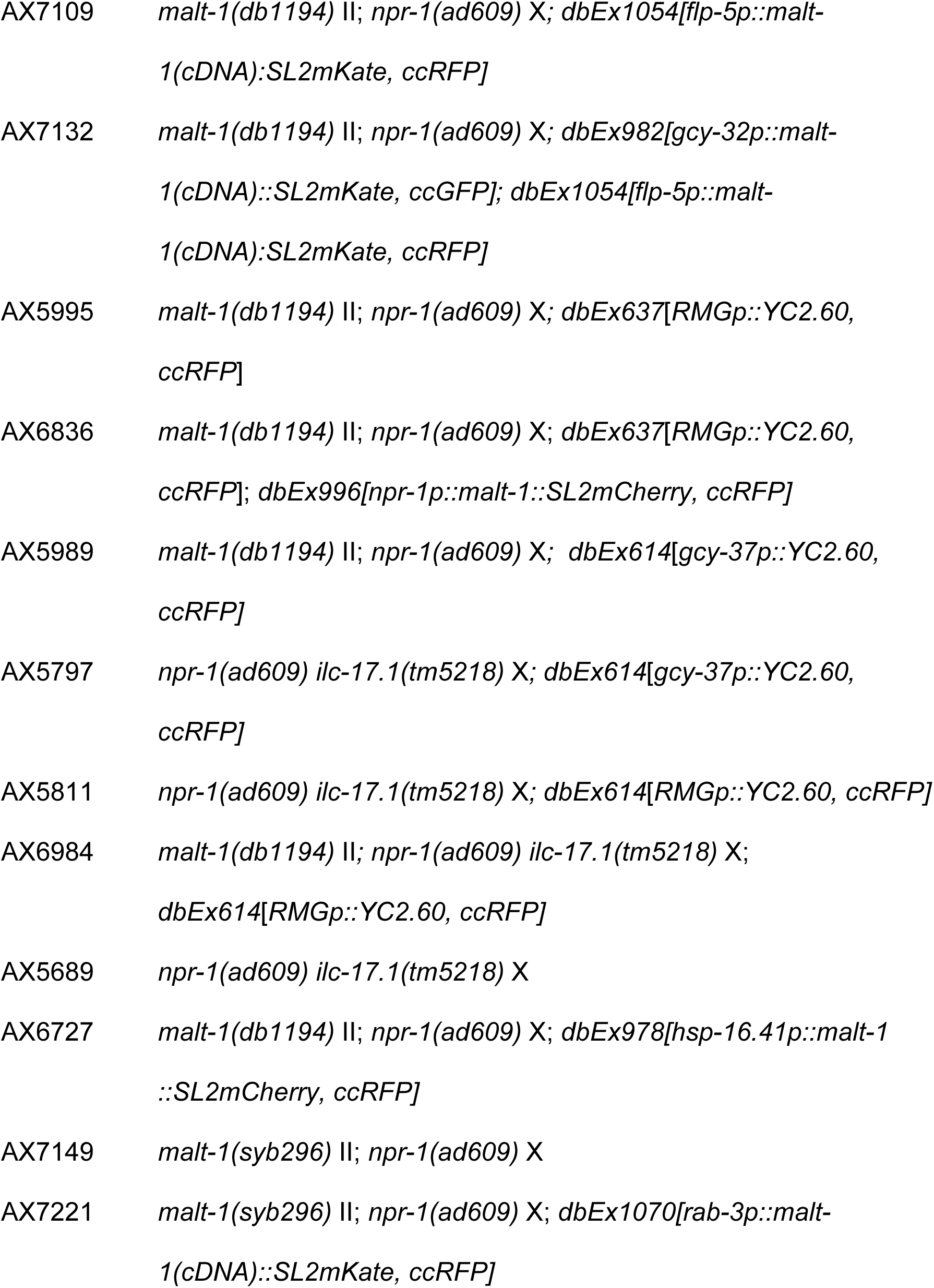
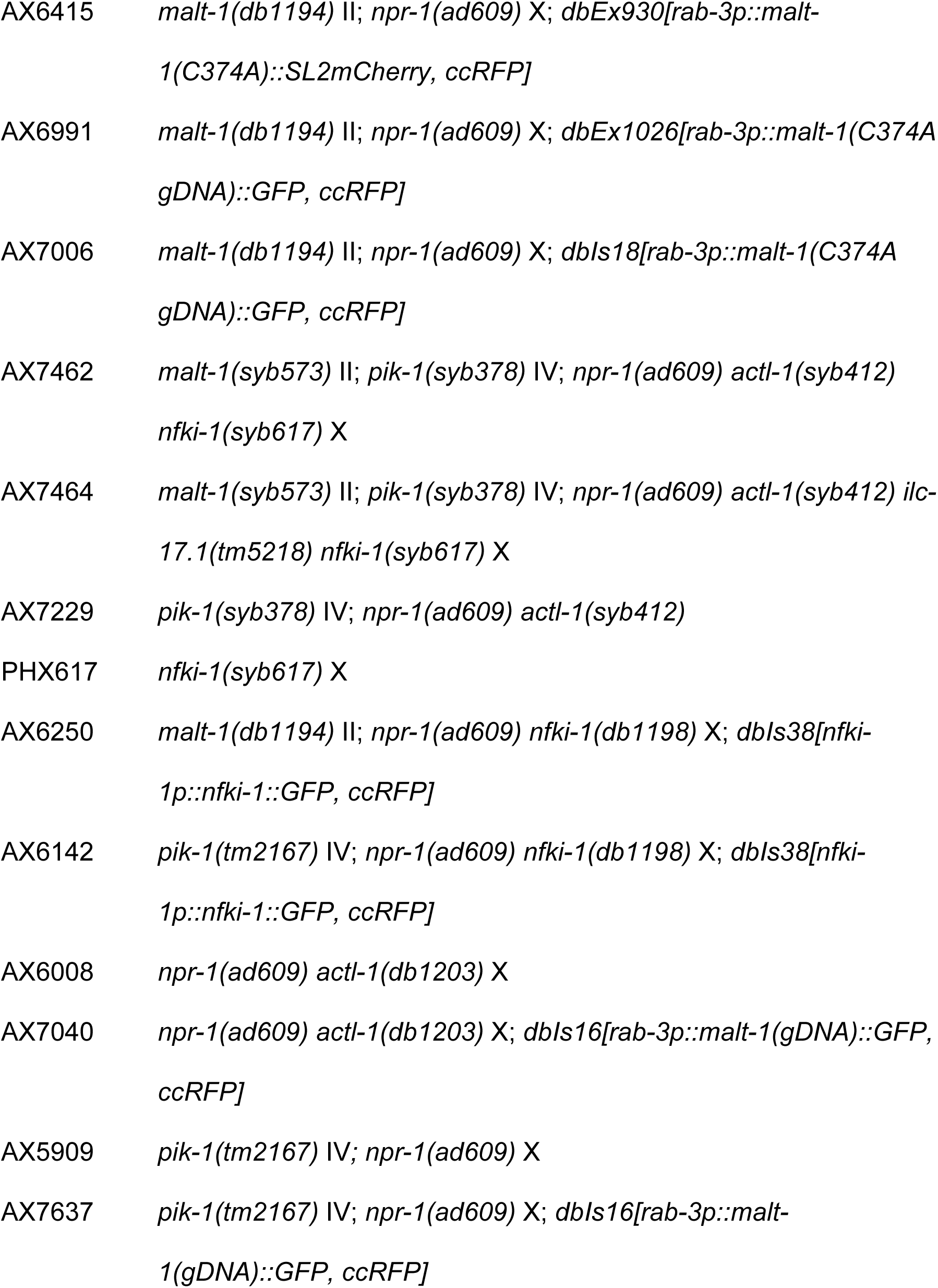
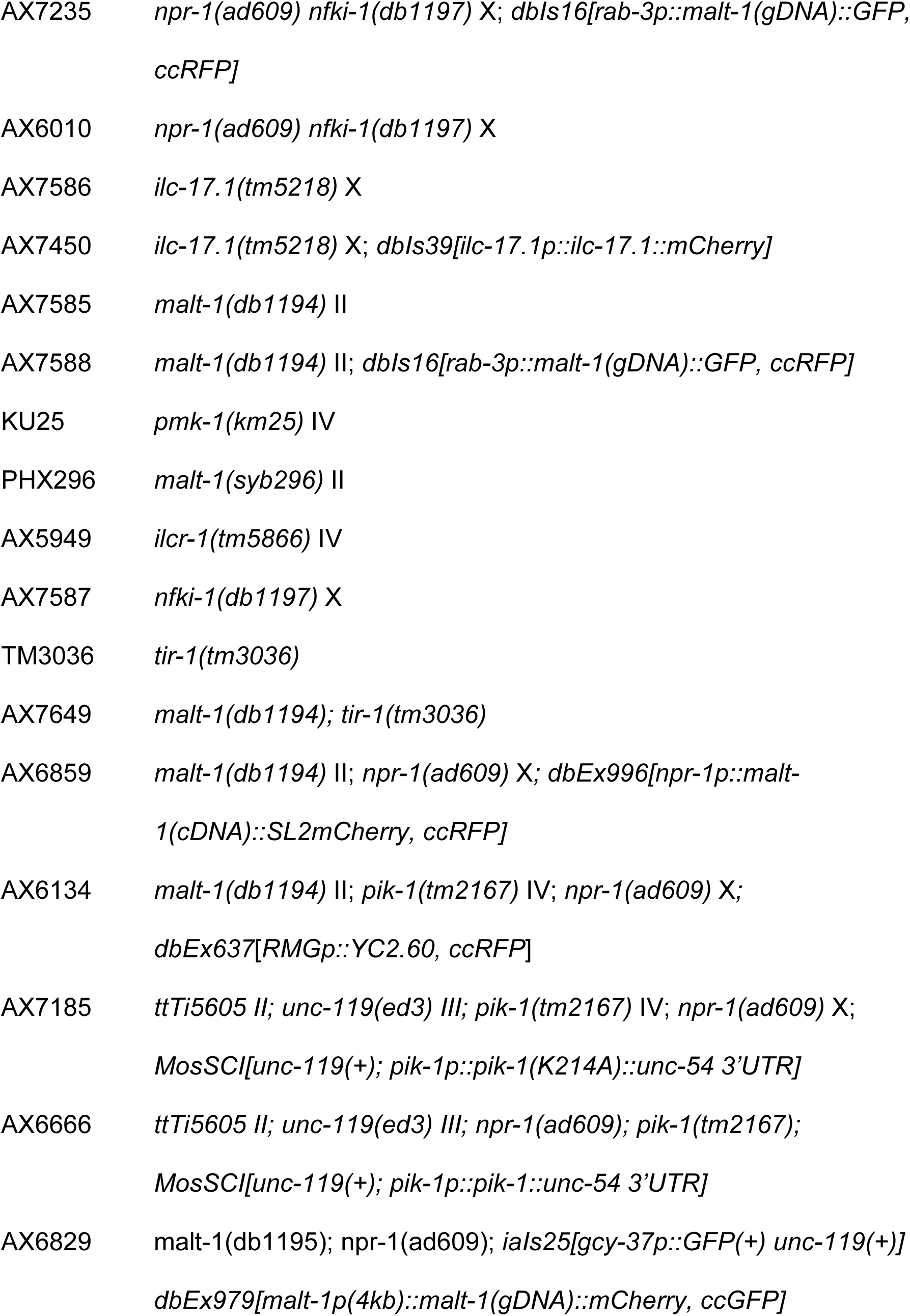
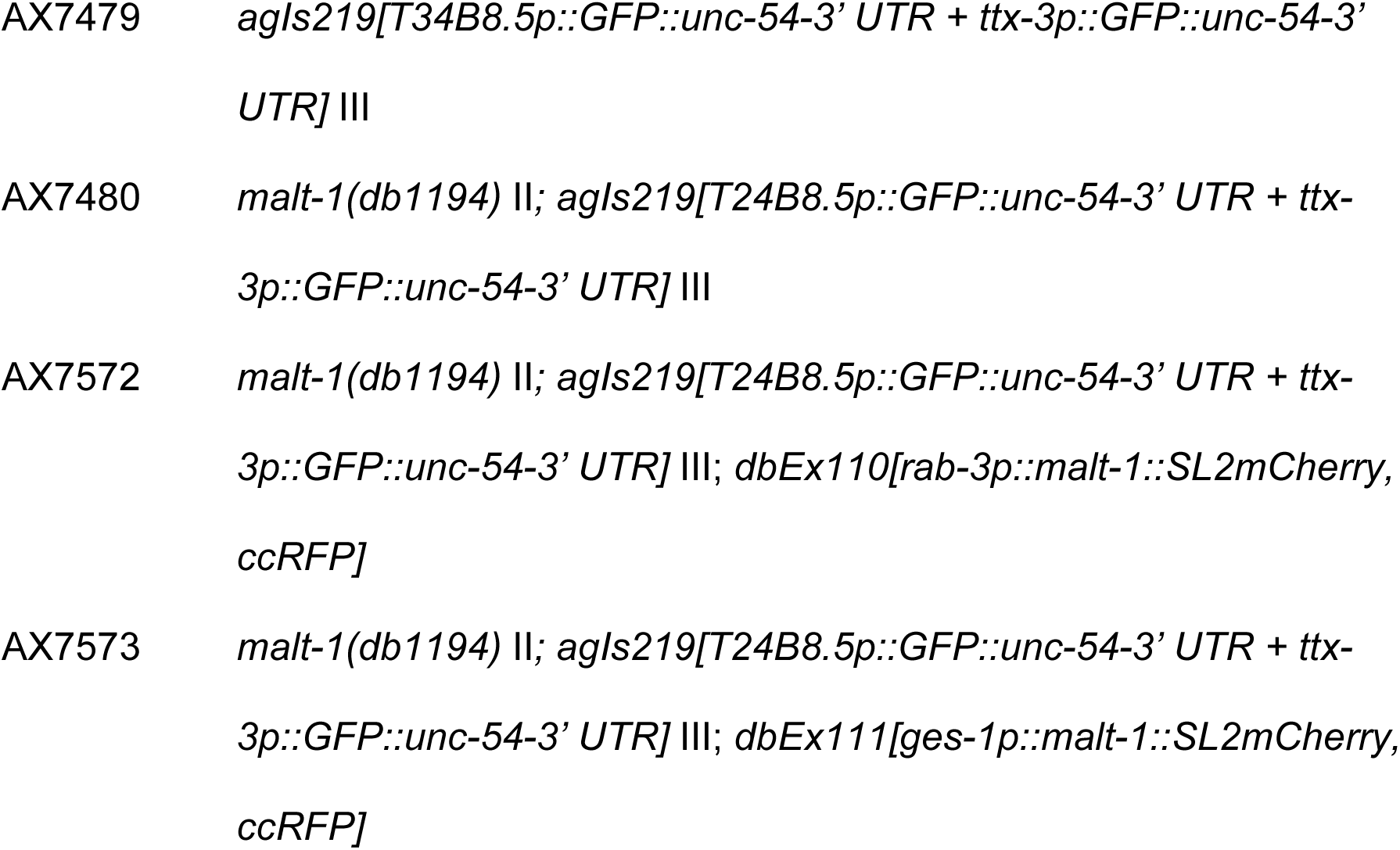
Strain list.

## Methods

### Strains and genetics

*C. elegans* were maintained on nematode growth medium (NGM) at room temperature (22°C) with *E. coli* OP50 food.

Whole genome sequencing showed that the aggregation-defective AX3621 strain was defective in *malt-1*. We used SNP mapping^66^ to investigate if the aggregation defect was linked to *malt-1*. We crossed AX3621 animals with the AX288 [*lon-2*(*e678*) *npr-1*(*ad609*)] strain; AX288 was constructed by backcrossing *lon-2 npr-1(ad609) X* 16x into the CB4856 (Hawaiian) wild strain. The *npr-1(ad609)* allele confers stronger aggregation than the CB4856 Hawaiian strain routinely used for mapping. We ‘singled’ 40-60 F2 animals, and scored their progeny for aggregation. Animals from non-aggregating F3 lines were pooled, and their DNA extracted and sequenced. Sequencing libraries were made using the Nextera DNA Library kit, and sequenced on a HiSeq 2500 (Illumina) machine with 125bp paired-end reads. Sequencing data was analyzed on using CloudMap^66^.

### Molecular Biology

*C. elegans* expression constructs were generated using MultiSite Gateway Recombination (Invitrogen). To amplify the *malt-1* promoter (4kb) we used primers ggggACAACTTTGTATAGAAAAGTTGctgc cggtggattccaacatattg and ggggACTGCTTTTTTGTACAAACTTGtctgaaattggggttcaagaaatttatttttgatttttaaaata, to amplify the *malt-1* ORF (gDNA): ggggACAAGTTTGTACAAAAAAGCAGGCTtttcagaaaaatgaacacaaacttggcggagttc and ggggACCACTTTGTACAAGAAAGCTGGGTActgtagacatttgattcttgtaatcaa aatatgaccaatatc and to amplify *malt-1* cDNA: ggggACAAGTTTGTACAAAAAAGCAGGCTtttcagaaaaatgaacacaaacttggcggagttc and ggggACCACTTTGTACAAGAAAGCTGGGTATTACTGTAGACATTTGATTCTTGTAATCAAAATATGACCAATATCAACATTC.

The Q5 Site-Directed Mutagenesis Kit (NEB) was used to create *malt-1(C374A)* cDNA, with the following primers: TCTTGATGTCgcCAGAAAATTTGTTCCATATG and gcgcgtcaagttgtGCCTGACGACGAGTTGTGCTGTTTTAGAGCTAGAA.

To generate deletions in the *malt-1* locus by CRISPR/Cas9 we expressed a gaucagguauccaccguag short guide from the *rpr-1* promoter, as described^67^. The primers used to amplify this sequence for insertion into an *E*coRI-cut expression plasmid (addgene #48961) were gcgcgtcaagttgtGgatcaggtatccaccgtagGTTTTAGAGCTAGAA and TTCTAGCTCTAAAACctacggtggatacctgatcCacaacttgacgcgc.

Expression constructs were injected at 50 ng/µl, with the exception of CRISPR-Cas9 mixes that were prepared as previously described^67^: 30ng ng/µl *eft-3::cas9*, 100 ng/µl sgRNA, 30ng ng/µl cc::GFP.

The following alleles were generated by SunyBiotech (Fuzhou, China) using CRISPR/Cas9-based genome editing: *malt-1(syb296)*, *actl-1(syb412), pik-1(syb378), malt-1(syb573)*, and *nfki-1(syb617)*. We verified modified sequences using Sanger sequencing (Supplementary Table 6).

### Behavioral assays

Behavioral assays were performed at room temperature (22°C). Aggregation was assayed as previously described^45^. Briefly, 60 young adults were picked onto a plate seeded with 100 µl OP50 48h previously. Animals were left undisturbed for 2h and then scored blind to genotype. The % of animals in groups was calculated, with a group defined as 3 or more animals in contact with one another.

Locomotory responses to O_2_ stimuli were measured as described previously^29, 68^ with minor modifications. 15-25 young adults were picked onto a plate seeded with 20 µl OP50 14-18h previously, and covered with a microfluidic PDMS chamber. Defined O_2_ mixtures (balance nitrogen) were bubbled through H_2_O and delivered to the PDMS chamber at a rate of 1.4ml/min using a PHD 2000 Infusion syringe pump (Harvard Apparatus). Video recordings were acquired at 2fps with FlyCapture software, using a Point Grey Grasshopper camera mounted on a Leica MZ6 microscope. Speed and reversals were measured using Zentracker custom software (https://github.com/wormtracker/zentracker). To measure phenotypes associated with IL-17 signaling defects, worms were left undisturbed for 2h on assay plates prior to recording.

To measure thrashing, single animals were placed into individual wells containing 50 µl M9 buffer. The number of complete body bends per minute was measured by a scorer who was blind to genotype.

### Heat-shock

As reported previously, the *hsp-16.41* heat shock promoter is leaky in animals grown at room temperature. We therefore kept animals at 15°C until the time of heatshock (late L4). To induce heat-shock, parafilm-wrapped plates were submerged in a 34°C water bath for 30 min, and then recovered at room temperature until the time of assay.

### Light microscopy

Worms were immobilized with 25 mM sodium azide on 2% agarose pads. Z stacks from animals expressing MALT-1::GFP and MALT-1::RFP were acquired on an Inverted Leica SP8 confocal microscope using a 63x/1.20 water objective. Figure panels were obtained using the Z-project (average intensity) function in Image J.

We quantified GFP intensity in L4 animals expressing the *agIs219*(*pT24B8.5::GFP*) transgene^48^ using a Nikon Ti2 microscope with a Niji LED light source (Bluebox Optics, Huntingdon, UK and a NEO scientific CMOS camera (Andor, Belfast, UK), with a 10x objective (Nikon, Tokyo, Japan) and 50 ms exposure time.

### Calcium Imaging

Animals expressing cameleon YC2.60 were imaged with a 2x AZ-Plan Fluor objective (Nikon) on a Nikon AZ100 microscope fitted with ORCA-Flash4.0 digital cameras (Hamamatsu). Excitation light was provided from an Intensilight C-HGFI (Nikon), through a 438/24nm filter and an FF458DiO2 dichroic (Semrock). Emission light was split using a TwinCam dual camera adaptor (Cairn Research) and passed through CFP (483/32nm) and YFP (542/27) filters, and a DC/T510LPXRXTUf2 dichroic. We acquired movies using NIS-Elements, with 100 ms exposure time.

To image neural activity in freely moving animals (Supplementary Fig. 4g), single young adults were transferred to peptone-free agar plates spotted with 4 µl of concentrated OP50 food in M9 buffer, and imaged at 2x zoom. For all other figures, 4-8 young adults were transferred to peptone-free agar plates, immobilized on a 2µl patch of concentrated OP50 in M9 buffer using Dermabond adhesive, leaving the nose exposed, and imaged at 4x zoom.

### Immunoprecipitation from *C. elegans*

For co-IP experiments analyzed by LC-MS/MS, *C. elegans* lysis and affinity purification was performed as previously described^69^ with minor modification. Lysis buffer A was prepared with 50 mM HEPES (pH 7.4), 1 mM EGTA, 1 mM MgCl_2_, 100 mM KCl, 10% glycerol, 0.05% NP40, 1mM DTT, 0.1 M PMSF and 1 complete EDTA-free proteinase inhibitor cocktail tablet (Roche Applied Science) per 12 ml. Unsynchronized worms grown in liquid were washed twice in M9 and once in ice-cold lysis buffer A, then snap-frozen by dropwise addition to LN_2_ in preparation for cryogenic grinding. Worm “popcorn” was pulverized using a Freezer/Mill (SPEX SamplePrep). Crude extract was clarified at 4°C for 10 min at 20,000*g,* and again for 20 min at 100,000*g* with a TLA-100 rotor (Beckman Coulter). For IP, roughly equal volumes of sample and control lysate were incubated with 100 µl GFP-Trap MA (ChromoTek gtma), Myc-Trap MA (ChromoTek ytma), or anti-FLAG M2 magnetic beads (Sigma M8823) for 3-4h at 4°C, then washed twice with 50mM HEPES, 100mM KCl. Purified complexes were eluted in SDS-sample buffer at 95°C and fractionated by SDS-PAGE prior to characterization by LC-MS/MS.

For co-IP experiements analysed by Western blot, the following modifications were made. Lysis buffer B contained 50 mM HEPES (pH 7.4), 100 mM KCl, 0.05% NP40, 1 mM DTT, 0.1 M PMSF and 1 complete EDTA-free proteinase inhibitor cocktail tablet (Roche Applied Science) per 12ml. Crude extract was clarifired at 4°C for 30 min at 14,000 rpm. For immunoprecipitation, samples were incubated with anti-HA agarose (Sigma A2095), anti-HA magnetic beads (Cell Signaling #11846), anti-FLAG M2 magnetic beads, or anti-V5 agarose (Sigma A7435) for 3-4h at 4°C, then washed 5x with 50 mM HEPES, 100 mM KCl.

### Identification of protein-protein interactions by MS

Gel samples were destained with 50% v/v acetonitrile and 50 mM ammonium bicarbonate, reduced with 10 mM DTT, and alkylated with 55 mM iodoacetamide. Proteins were digested with 6 ng/µl trypsin (Promega, UK) overnight at 37°C, and peptides extracted in 2% v/v formic acid 2% v/v acetonitrile, and analysed by nano-scale capillary LC-MS/MS (Ultimate U3000 HPLC, Thermo Scientific Dionex) at a flow of ∼ 300 nL/min. A C18 Acclaim PepMap100 5 µm, 100 µm × 20 mm nanoViper (Thermo Scientific Dionex), trapped the peptides prior to separation on a C18 Acclaim PepMap100 3 µm, 75 µm × 250 mm nanoViper. Peptides were eluted with an acetonitrile gradient. The analytical column outlet was interfaced via a nano-flow electrospray ionisation source with a linear ion trap mass spectrometer (Orbitrap Velos, Thermo Scientific). Data dependent analysis was performed using a resolution of 30,000 for the full MS spectrum, followed by ten MS/MS spectra in the linear ion trap. MS spectra were collected over a m/z range of 300–2000. MS/MS scans were collected using a threshold energy of 35 for collision-induced dissociation. LC-MS/MS data were searched against the UniProt KB database using Mascot (Matrix Science), with a precursor tolerance of 10 ppm and a fragment ion mass tolerance of 0.8 Da. Two missed enzyme cleavages and variable modifications for oxidised methionine, carbamidomethyl cysteine, pyroglutamic acid, phosphorylated serine, threonine and tyrosine were included. MS/MS data were validated using the Scaffold programme (Proteome Software Inc).

### Quantification of NFKI-1 interacting peptides by TMT labeling

#### On-beads trypsin digestion

Protein samples on beads were reduced with 10 mM DTT at 56°C for 30 min and alkylated with 15 mM iodoacetamide (IAA) in the dark at 22°C for 30 min. Alkylation was quenched by adding DTT and the samples digested with trypsin (Promega, 1.25µg) overnight at 37°C. After digestion, supernatants were transferred to a fresh Eppendorf tube, the beads were extracted once with 80% acetonitrile/ 0.1% TFA and combined with the corresponding supernatant. The peptide mixtures were then partially dried in a Speed Vac and desalted using home-made C18 (3M Empore) stage tip filled with 4 µl of poros R3 (Applied Biosystems) resin. Bound peptides were eluted sequentially with 30%, 50% and 80% acetonitrile in 0.1%TFA and lyophilized.

#### Tandem mass tag (TMT) labeling

Dried peptide mixtures from each condition were re-suspended in 40µl of 250mM triethyl ammonium bicarbonate. 0.8mg of TMT 6plex reagents (Thermo Fisher Scientific) was re-constituted in 41µl anhydrous MeCN. 20µl of TMT reagent was added to each peptide mixture and incubated for 1 hr at r.t. The labeling reactions were terminated by incubation with 4.4µl of 5% hydroxylamine for 15min. For each condition, the labeled samples were pooled, Speed Vac to remove acetonitrile, desalted and then fractionated with home-made C18stage tip using 10mM ammonium bicarbonate and acetonitrile gradients. Eluted fractions were acidified, partially dried down in speed vac and ready for LC-MSMS.

#### LC-MSMS

Peptides were separated on an Ultimate 3000 RSLC nano System (Thermo Scientific), using a binary gradient consisting of buffer A (2% MeCN, 0.1% formic acid) and buffer B (80% MeCN, 0.1% formic acid). Eluted peptides were introduced directly via a nanospray ion source into a Q Exactive Plus hybrid quardrupole-Orbitrap mass spectrometer (Thermo Fisher Scientific). The mass spectrometer was operated in standard data dependent mode, performed survey full-scan (MS, m/z = 350-1600) with a resolution of 140000, followed by MS2 acquisitions of the 15 most intense ions with a resolution of 35000 and NCE of 33%. MS target values of 3e6 and MS2 target values of 1e5 were used. Dynamic exclusion was enabled for 40s.

#### Data analysis – MaxQuant

The acquired MSMS raw files were processed using MaxQuant (Cox and Mann) with the integrated Andromeda search engine (v.1.5.5.1). MSMS spectra were searched against the *Caenorhabditis elegans* UniProt Fasta database (July 2017). Carbamidomethylation of cysteines was set as fixed modification, while methionine oxidation and N-terminal acetylation (protein) were set as variable modifications. Protein quantification required 1 (unique+ razor) peptide. Other parameters in MaxQuant were set to default values. MaxQuant output file, proteinGroups.txt was then processed with Perseus software (v 1.5.5.0). After uploading the matrix, the data was filtered, to remove identifications from reverse database, modified peptide only, and common contaminants. Each peptide channel was normalized to the median and log2 transformed.

### Sub-cellular fractionation

Nuclear/cytoplasmic fractionation was perfomed as described previously^70^. Briefly, young adult worms were washed 3-5 times in M9, and twice in hypotonic buffer (15mM HEPES, 10mM KCl, 5mM MgCl_2_, 0.1mM EDTA, 350mM sucrose). Lysis was on ice in complete hypotonic buffer plus 1mM DTT and 1 complete EDTA-free proteinase inhibitor cocktail tablet (Roche Applied Science) per 12ml, using a motorized pellet pestle (Sigma Z359971, Z359947) until most worm carcases were homogenized. Worm debris was pelleted at 500*g* (2 × 5min), and 5% of the resultant supernatant was kept as the input fraction. Nuclei were pelleted at 4,000*g* (5 min), and the resulting supernatant centrifuged again at 17,000*g* and kept as the cytoplasmic fraction. Nuclear pellets were washed twice in complete hypotonic buffer and dissolved in complete hypertonic buffer (15mM HEPES, 400mM KCl, 5mM MgCl_2_, 0.1mM EDTA, 0.1% Tween 20, 10% glycerol, 1mM DTT, complete EDTA-free proteinase inhibitor as above).

### Size-exclusion chromatography

*C. elegans* lysate was prepared as described above for LC-MS/MS, and loaded onto a Superose 6 Increase 10/300 GL column. The column was equilibrated with Lysis buffer A and 1ml fractions were collected.

### Immunoblotting

After SDS-PAGE using Bolt 4-12% Bis-Tris Plus gels (ThermoFisher Scientific), protein was transferred to PVDF membrane (0.45-micron pore size, ThermoFisher Scientific) using the XCell II Blot Module (ThermoFisher Scientific). Membranes were blocked with 5% milk for 1h, then incubated with primary antibody overnight at 4°C, followed by secondary antibody for 1h at RT. Unbound antibody was washed away with PBST (3 × 5min), and SuperSignal West Pico PLUS Cemiluminescent Substrate (ThermoFisher Scientific) used for detection. The following commercially available antibodies were used: anti-FLAG M2-Peroxidase (Sigma A8592), anti-Myc (Cell Signaling #2276), anti-HA (Cell Signaling #3956), anti-V5 (Bethyl Laboratories A190-120A), anti-phospho-p38 MAPK (Cell Signaling #4511), anti-Histone H3 (Cell Signaling #9715), anti-alpha tubulin (abcam ab40742). phospho-PMK-1 western blot assays were performed as described in^71^.

### RNA-seq

#### RNA preparation

10 Gravid adults were left to lay eggs for 2h on an OP50 lawn seeded 24h previously, before being picked away. 8-10 plates were used per replicate, and all genetics backgrounds were prepared in parallel. Once animals reached late L4 stage, they were washed twice in M9 and frozen in LN_2_. 1ml Qiazol Lysis Regaent (Qiagen) and 300-400µl 0.7mm Zirconia beads (BioSpec) were added to worm pellets in preparation for mechanical disruption. Samples were disrupted with a TissueLyser (Qiagen), using 1min at maximum power followed by 1min on ice (repeated 4 times). RNA was extracted using the RNeasy Plus Universal Mini Kit (Qiagen), following the manufacturer’s instructions with the exception that 1-Bromo-3-chloropropane (Sigma, B673) was used instead of chloroform.

#### Library preparation, sequencing and analysis

Libraries were prepared using the TruSeq Stranded mRNA kit (Illumina) with polyA capture for mRNA, and sequenced on a HiSeq 4000 platform (Illumina) with 50 bp single-end reads. We sequenced five independent biological replicates for *npr-1 ilc-17.1*, and six for *npr-1*, *malt-1; npr-1*, and *npr-1 nfki-1.* Reads were aligned to the *C. elegans* genome using TopHat v2.1.0, and expression quantified using Cufflinks v2.2.1. For statistical comparisons a *q-* value <0.05 was considered significant. Only genes for which the FPKM (Fragments Per Kilobase of transcript per Million mapped reads) was ≥ 1 in *npr-1*, and for which the log2(fold change) between conditions was ≥0.25 were included in Fig. 6. Genes whose expression oscillates during development^72^ were excluded.

#### GO enrichment

GO terms for these genes were retrieved and GO enrichment calculated using g:Profiler (version e94_eg41_p11_592d917^65^). Terms with multiple testing corrected p-values < 0.05 were considered enriched.

### Associative learning assay

Chemotaxis assays were performed as previously described^73^ with minor modification. To establish salt gradients 100mM NaCl agar plugs were left overnight on assay plates containing 1mM CaCl_2_, 1mM MgSO_4_, 25mM K_2_HPO_4_ pH 6. Plugs were removed immediately before the assay, and replaced with 1µl 1M NaN_3_. Another 1µl NaN_3_ was added equidistant from the starting point as a control. The chemotaxis index was calculated as (A – B)/N, where A was the number of animals within 1 cm of the peak of the salt gradient, B was the number of animals within 1 cm of the control spot, and N was the number of all animals. Conditioning was performed as described previously^49^. Briefly, synchronized young adults raised on OP50 were washed three times in CTX buffer (5 mM K_2_HPO_4_ pH 6, 1 mM CaCl_2_ and 1 mM MgSO_4_), then left for 4h on NGM agar with no NaCl (mock) or 300mM NaCl (conditioned).

### *P. aeruginosa* killing assays

Slow killing assays were performed as described^71^ with 10µM 5-fluorodeoxyuridine (FUdR). Synchronized L4s raised on OP50 were added to 0.35% peptone NGM plates, seeded the day before with PA14. Animals were scored every 12h, and counted as dead if they did not respond to prodding. Logrank tests with Bonferroni correction were performed using OASIS (On- line Application for Survival Analysis, http://sbi.postech.ac.kr/oasis^74^).

### Lifespan analyses

Life span was measured as described^75^, starting at day 1 of adulthood. Scoring and statistical analyses were performed as described above for *P. aeruginosa* killing assays.

#### Statistics

Statistical tests and n values used for experiments in this paper are indicated in the corresponding figure legend or methods section. For salt chemotaxis and aggregation behavior, statistical significance between groups was determined using one-way ANOVA with Tukey’s post hoc HSD. Differences in locomotory response and FRET levels (yellow cameleon) were evaluated by Mann-Whitney *U* test. Logrank tests with Bonferroni correction were used to compare genotypes in lifespan and PA14 survival assays.

